# Right inferior frontal cortex damage impairs the initiation of inhibitory control, but not its implementation

**DOI:** 10.1101/2022.05.03.490498

**Authors:** Yoojeong Choo, Dora Matzke, Mark D. Bowren, Daniel Tranel, Jan R. Wessel

**Author notes:** Corresponding author address: Jan R. Wessel, Ph.D. Department of Neurology, University of Iowa Hospitals and Clinics 200 Hawkins Drive, Iowa City IA 52242, wessellab.org.

## Abstract

Inhibitory control is one of the most important control functions in the human brain. Much of our understanding of its neural basis comes from seminal work showing that lesions to the right inferior frontal cortex (rIFC) increase stop-signal reaction time (SSRT), a latent variable that expresses the speed of inhibitory control. However, recent work has identified substantial limitations of the SSRT method. Notably, SSRT is confounded by trigger failures: stop-signal trials in which inhibitory control was never initiated. Such trials inflate SSRT, but are typically indicative of attentional, rather than inhibitory deficits. Here, we used hierarchical Bayesian modeling to identify stop-signal trigger failures in human rIFC lesion patients, non-rIFC lesion patients, and healthy comparisons. Furthermore, we measured scalp-EEG to detect β-bursts, a neurophysiological index of inhibitory control. rIFC lesion patients showed a more than five-fold increase in trigger failure trials and did not exhibit the typical increase of stop-related frontal β- bursts. However, on trials in which such β-bursts did occur, rIFC patients showed the typical subsequent upregulation of β over sensorimotor areas, indicating that their ability to implement inhibitory control, once triggered, is intact. These findings suggest that the role of rIFC in inhibitory control has to be fundamentally reinterpreted.

## Introduction

Humans have remarkable cognitive control abilities, which allow them to safely navigate complex everyday situations. For example, when crossing a street, humans can rapidly stop themselves from continuing to walk when they suddenly notice a rapidly approaching car. The process underlying this ability to stop an already-initiated action is inhibitory control. In the laboratory, inhibitory control is typically tested in the stop-signal task (Verbruggen et al., 2019), in which an initial go-signal (i.e., a cue to initiate a movement) is sometimes followed by a subsequent stop-signal, prompting the cancellation of that movement. The processes underlying behavior in the stop-signal task are described in a well-characterized cognitive model – the horse-race model (Logan & Cowan, 1984). This model purports that on each trial, the inhibitory process triggered by the stop-signal races with the movement-initiation process triggered by the go-signal, thereby determining whether an action can be successfully stopped. The assumptions of the horse-race model allow the calculation of stop-signal reaction time (SSRT) – a latent variable that expresses the speed of the stopping process, which is not otherwise overtly observable (as successful stopping is defined by the absence of a response). In a seminal study on the neural basis of inhibitory control, Aron and colleagues (2003) showed that lesions to the right inferior frontal cortex (rIFC) cause an elongation of SSRT, prompting the proposal that “response inhibition can be localized to a discrete region of the PFC” – namely, the rIFC. This finding has spawned a wider, highly-influential theory of inhibitory control, which, in its most recent iteration, holds that “rIFC implements a brake over response tendencies” (Aron et al., 2014).

However, the SSRT method underlying this (and other) seminal work has recently undergone several substantial challenges (e.g., Bissett et al., 2021; Matzke, Love, et al., 2017). One of the most prominent shortcomings of SSRT is that it does not account for trigger failures (Band et al., 2003) – trials with stop-signals in which inhibitory control process was never initiated to begin with. On such trials, erroneously executed responses do not result from an insufficiently fast stop-process losing the horse race, but from the mere fact that inhibitory control never ‘entered the race’. Crucially, including such trials in the SSRT calculation leads to artificially inflated SSRT estimates and – in the worst-case – can produce fictitious group differences in inhibitory control speed, which are instead more likely due to attentional lapses (e.g., Matzke, Hughes, et al., 2017). In other words, in the above-mentioned example of crossing a street, a trigger failure could be indicative of a deficit in detecting an approaching car, rather than in intercepting the movement fast enough.

In light of this confound, it is necessary to reassess the causal link between stop-signal performance and rIFC damage. Therefore, we here repeated Aron et al. (2003)’s original lesion study with a stop-signal task that was optimized to implement a hierarchical Bayesian technique for the identification of trigger failures (Matzke, Love, et al., 2017). We specifically tested whether rIFC lesion patients show increased trigger failure rates, as rIFC is a key region that is often implicated in stimulus-driven attention more generally (Corbetta & Shulman, 2002). Moreover, we extended the behavioral investigation by Aron and colleagues by measuring scalp-EEG. Specifically, we aimed to investigate the influence of rIFC lesions on stop-related β-bursts dynamics. β-bursts are a recently discovered neurophysiological signature of inhibitory control (Diesburg et al., 2021; Wessel, 2020) and can provide additional insights into the distinction between the initiation and the implementation of inhibitory control. Specifically, β-burst rates over frontal cortex are increased on stop- compared to go-trials (Enz et al., 2021; Wessel, 2020; Jana et al., 2020), which has been proposed to reflect the initial stage of the inhibitory cascade that ultimately results in action-stopping. In other words, frontal β-bursts reflect the *initiation* of inhibitory control. At the other end of the cascade, β-bursts over sensorimotor areas can be used to measure the successful *implementation* of inhibitory control. Sensorimotor β activity reflects an inhibited state of the motor system at baseline (Kilavik et al., 2013; Soh et al., 2021), and stop-related frontal β-bursts are followed by a rapid re-instantiation of these sensorimotor bursts (Wessel, 2020). This purportedly reflects the successful implementation of the inhibitory cascade and a return to an inhibited motor system (Diesburg et al., 2021).

Hence, in line with our behavioral hypothesis, we predicted that if rIFC lesion patients showed increased trigger failure rates, they would also show reduced frontal β-burst rates compared to both non-rIFC patients and healthy adult comparisons, reflecting a deficit in triggering/initiating inhibitory control. However, we furthermore predicted that when a frontal β- burst does take place in rIFC patients (i.e., when the cascade is successfully triggered), sensorimotor β-burst rates would be appropriately upregulated, reflecting a retained ability to implement inhibitory control, despite damage to rIFC.

## Results

### Participants

Participants included 16 rIFC lesion patients, 16 non-rIFC lesion patients, and 32 age- and sex-matched comparisons. Lesion overlap maps are found in ***Figure 1*** and demographic data for all participants are presented in ***Table 1***. All participants performed a version of the stop-signal task (***Figure 2A***) that was optimized for the usage of BEESTS, a hierarchical Bayesian modeling technique that simultaneously accounts for the shape of Go-RT and SSRT distributions and the prevalence of trigger failures in the stop-signal task (Matzke et al., 2013; Matzke, Love, et al., 2017, ***Figure 2B***).

**Figure 1.**
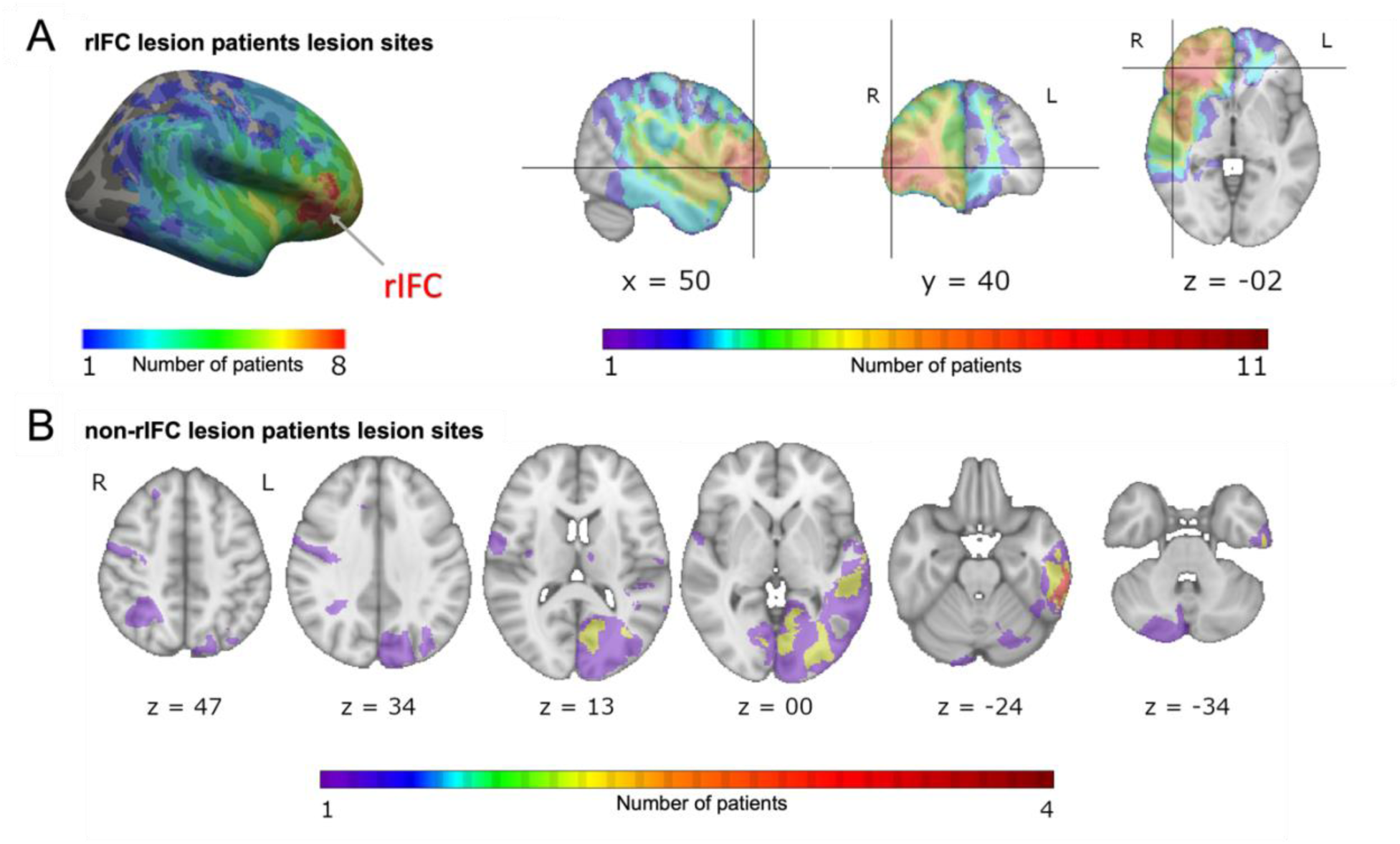
Overlap of lesions in patients with rIFC lesions (rIFC lesion patients, top, A) and in patients with lesions outside of rIFC (non-rIFC lesion patients, bottom, B). **A**, Top left, A lateral view of the lesion overlaps for rIFC lesion patients at the cortical level. Top right, (in order): sagittal, coronal, and axial view of the lesion overlap for rIFC lesion patients. The crosshair is centered on the right IFG (Inferior Frontal Gyrus) on the Harvard-Oxford Atlas. All rIFC patients were included (N = 16). Bottom, **B**, Axial view of the lesion overlap for non-rIFC lesion patients. One lesion mask for a patient was missing and not included (N = 15). The color bar indicates the number of patients overlapped in lesion sites. R: right, L: left.

**Table 1.** Demographic information all four groups

| Group | Sex | Handedness | Age | Chronicity |
| --- | --- | --- | --- | --- |
| rIFC lesions | 10M/6F | 16R/0L | 54.25 (15.19) | 26.88 |
| rIFC comparison | 10M/6F | 15R/1L | 54.63 (15.28) | n/a |
| Non-rIFC lesions | 10M/6F | 13R/3L | 61.50 (14.62) | 5.72 |
| Non-rIFC comparison | 10M/6F | 15R/1L | 61.94 (14.40) | n/a |

Abbreviations: M, male; F, female; R, right-handed; L, left-handed; Age: mean age at testing in years (standard deviation); Chronicity: median length of time between lesion onset and current experiment in years.

**Figure 2.**
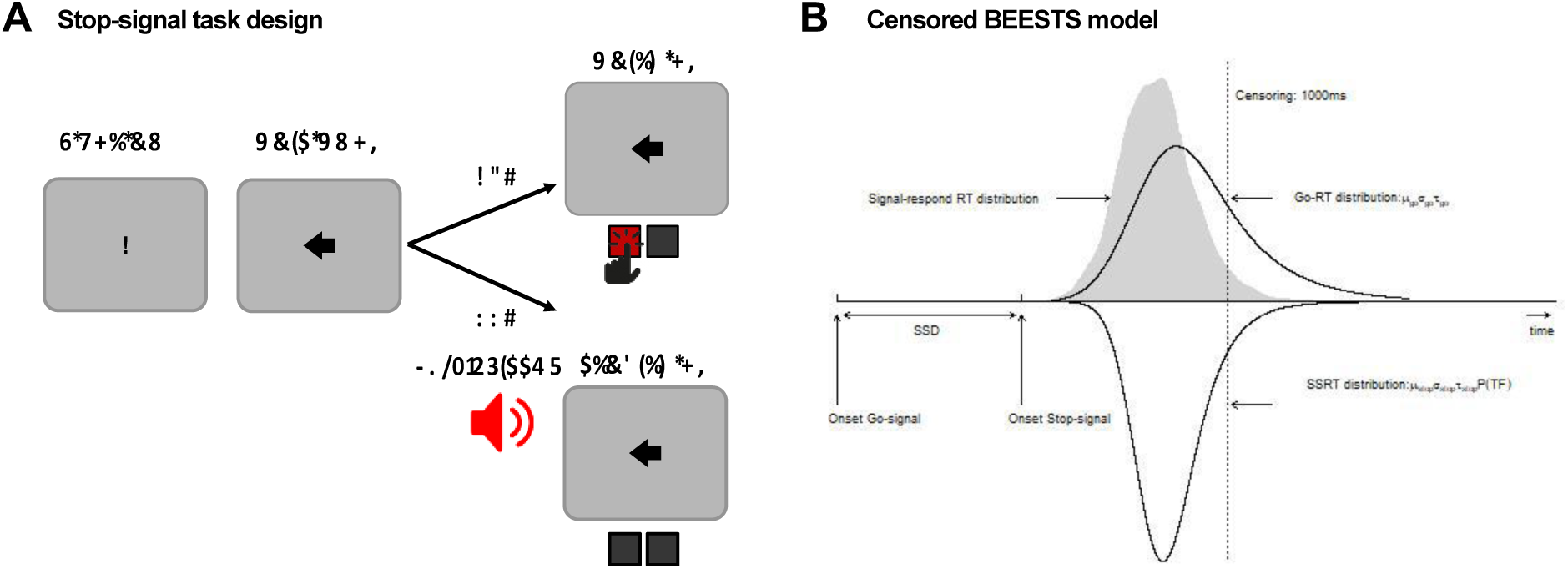
A) Schematic design of the stop-signal task. B) Graphical representation of the censored BEESTS model. The model assumes that the finishing times of the go (Go-RT distribution) and the stop runner (SSRT distribution) follow an ex-Gaussian distribution with parameters µgo, ***τ***_go_, and τgo, and µstop, σstop, and τstop, respectively. The finishing time distributions, and hence also the observed distribution of signal-respond RTs, is assumed to be censored above at 1000 ms to accommodate trials where the runners did not finish before the response window. The trigger failure parameter, P(TF), quantifies the probability that the stop runner was not initiated.

### Behavioral and modeling results

BEESTS assumes that the finishing times of the go (Go-RT distribution) and the stop runners (SSRT distribution) follow an ex-Gaussian distribution with parameters ***µ***, ***σ***, and ***τ***. The ***µ*** and ***σ*** parameters reflect the mean and the standard deviation of the Gaussian component and ***τ*** gives the mean of the exponential component and reflects the slow tail of the distribution. The mean and variance of the finishing time distributions can be obtained as ***µ*** + ***τ*** (i.e., mean Go-RT and SSRT) and ***σ*** ^2^ + ***τ*** ^2^, respectively. Using a mixture-likelihood approach, the model can be augmented with a parameter, P(*TF*), that quantifies the probability that participants fail to trigger the stop runner (Matzke, Love, et al., 2017). ***Table 2*** presents the posterior means and the corresponding 95% CIs of the population-level mean parameters in the four groups. The overlap of the posterior distributions of the go parameters indicated that ***μ***_go_ was lower whereas ***τ***_go_ was higher in the rIFC lesion group relative to matched comparisons, suggesting that the Go-RT distribution of the lesion group had a faster leading edge but a larger skew. Resulting from the nearly perfect trade-off between ***μ***_go_ and ***τ***_go_, mean Go-RT did not differ between the two groups. The overlap of the posterior distributions of the stop parameters indicated that ***μ***_stop_, ***σ***_stop_, and mean SSRT were higher in the rIFC lesion group than in matched comparisons, although the difference in ***σ***_stop_ was small. Crucially, we found a more than five-fold increase in the P(TF) parameter of the lesion group (16% vs. 3% in healthy comparisons). These results suggest that poor stop-signal performance associated with rIFC lesions is mainly attributable to increased trigger failure rate and a slowing of the inhibitory process as a result of a shift in the leading edge of the SSRT distribution. In contrast, with the exception of ***σ***_stop_, our analyses did not indicate the presence of differences in go or stop parameters between non-rIFC lesion patients and their matched comparisons, neither did we find evidence for a difference in the probability of trigger failures. Taken together, our results are mostly consistent with an attentional rather than inhibitory account of stop-signal deficits associated with rIFC lesions.

**Table 2.** Posterior means and 95% credible intervals (CI) of the population-level mean parameters in the four groups.

|  |  | Lesion patients |  | Matched comparisons |  | Difference |  |  |
| --- | --- | --- | --- | --- | --- | --- | --- | --- |
|  |  | Posterior mean | 95% CI | Posterior mean | 95% CI | Posterior mean | 95% CI | Bayesian *p* |
| **rIFC** | $\mu_{go}$ | 479 | [435,521] | 578 | [505,650] | -99 | [-181,-15] | **0.01** |
| | $\sigma_{go}$ | 75 | [60,94] | 90 | [73,107] | -15 | [-39,10] | 0.12 |
| | $\tau_{go}$ | 154 | [111,216] | 76 | [54,109] | 78 | [25,145] | **0** |
|  | Mean Go-RT | 633 | [572,708] | 654 | [576,730] | -21 | [-124,80] | 0.34 |
| | $\mu_{stop}$ | 223 | [197,249] | 198 | [182,215] | 25 | [-6,55] | 0.06 |
| | $\sigma_{stop}$ | 40 | [26,58] | 24 | [16,37] | 16 | [-3,37] | 0.05 |
| | $\tau_{stop}$ | 45 | [30,65] | 34 | [22,50] | 11 | [-12,35] | 0.17 |
|  | Mean SSRT | 268 | [237,298] | 232 | [211,254] | 35 | [-2,73] | 0.03 |
|  | P(TF) | 0.16 | [0.08,0.29] | 0.03 | [0,0.12] | 0.13 | [0.01,0.27] | **0.02** |
| **Non-rIFC** | $\mu_{go}$ | 547 | [504,589] | 575 | [506,646] | -28 | [-109,52] | 0.24 |
| | $\sigma_{go}$ | 98 | [84,111] | 104 | [85,122] | -6 | [-29,17] | 0.30 |
| | $\tau_{go}$ | 106 | [79,139] | 85 | [61,116] | 21 | [-19,62] | 0.15 |
|  | Mean Go-RT | 653 | [602,706] | 660 | [586,735] | -7 | [-97,83] | 0.43 |
| | $\mu_{stop}$ | 215 | [191,239] | 199 | [184,215] | 16 | [-11,43] | 0.12 |
| | $\sigma_{stop}$ | 54 | [24,104] | 24 | [14,36] | 31 | [0.11,80] | **0.02** |
| | $\tau_{stop}$ | 29 | [18,44] | 26 | [13,44] | 3 | [-18,22] | 0.36 |
|  | Mean SSRT | 244 | [218,271] | 225 | [204,249] | 19 | [-12,49] | 0.11 |
|  | P(TF) | 0.02 | [0,0.05] | 0.04 | [0.02,0.08] | -0.02 | [0.07,0.02] | 0.09 |

*Note.* The parameters of the Go-RT and SSRT distributions are presented on the ms scale. The P(TF) parameter is presented on the probability scale. The posterior distribution of “Difference” is computed by subtracting the posterior samples of the matched comparison group from the corresponding samples of the lesion group; i.e., positive values indicate that the parameter is higher in the lesion group than in the matched comparison group. “Bayesian *p*” is computed as the proportion of posterior samples in the posterior distribution of the lesion group that is larger than in the matched comparisons group. Bayesian *p* values are calculated as $p = \min(p, 1-p)$ , i.e., as non-directional tests, with significant results at a two-sided *p* of .05 (i.e., a one-sided critical *p* of .025) highlighted in bold.

### EEG results: Frontal β-bursts

***Figure 3*** shows the results of the analysis of frontal β-bursts. The 2x3 ANOVA of frontal β-burst rates for the rIFC group and their matched comparisons revealed a main effect of TRIAL TYPE (*F*(2, 60) = 3.749, *p* = .029, *η*^2^ = .111) and, crucially, a significant interaction between GROUP and TRIAL TYPE (*F*(2, 60) = 4.215, *p* = .019, *η*^2^ = .123). The main effect of GROUP was not significant (*F*(1, 30) = .046, *p* = .832, *η*^2^ *=* .002).

**Figure 3.**
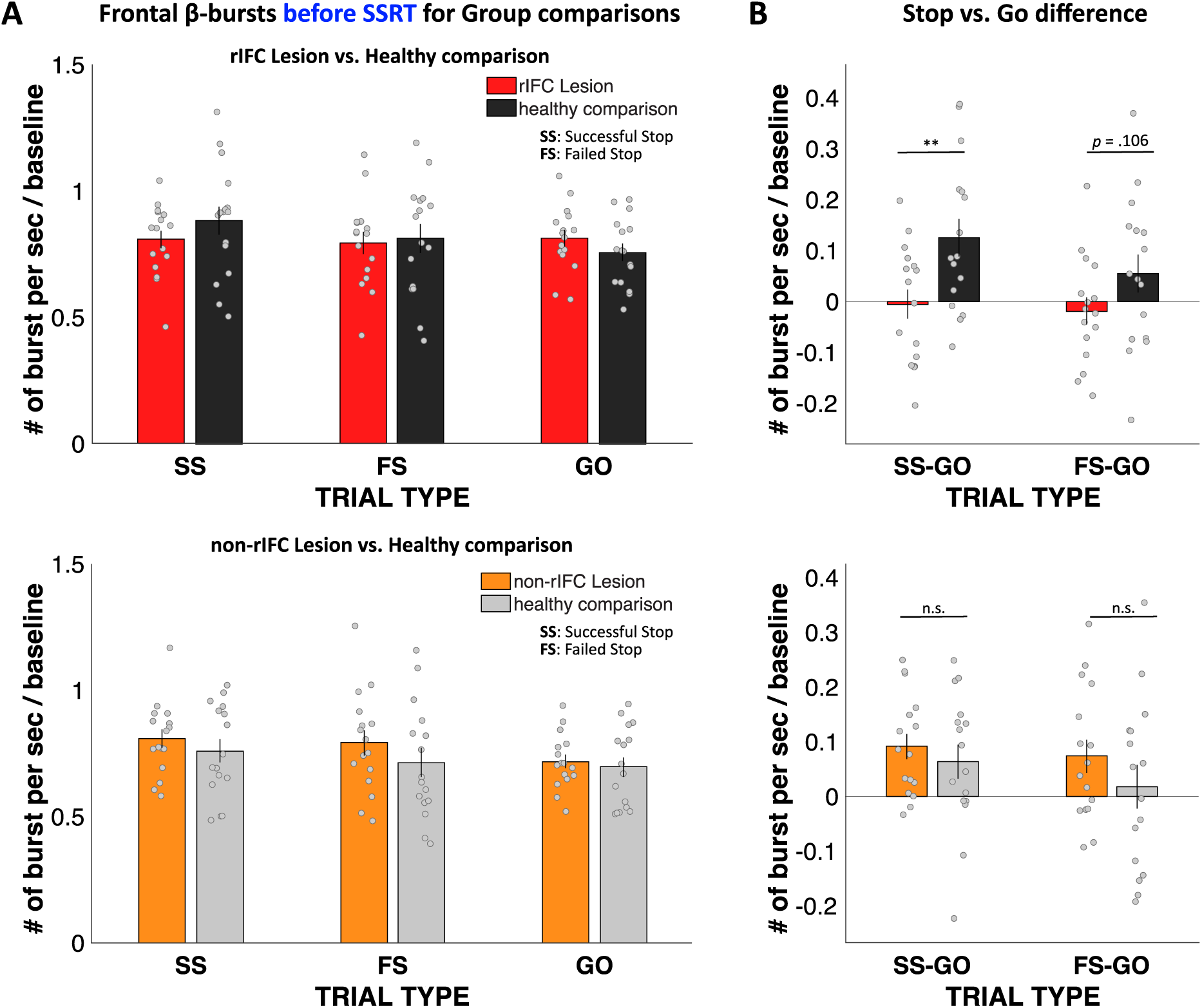
Normalized β-burst rate (per second) for group comparisons. **A**, Frontal β-burst rate before SSRT was shown for TRIAL TYPE (Successful Stop, Failed Stop, and Go) for two groups, respectively. Top: rIFC lesion group vs. matched healthy comparison group. Bottom: non-rIFC lesion group vs. matched healthy comparison group. **B,** Stop vs. Go difference (Successful Stop - Go and Failed Stop – Go). Top: rIFC lesion group vs. matched healthy comparison group. Bottom: non-rIFC lesion group vs matched healthy comparison group. Dots represent individual participant means. Error bar indicates ±SEM.

Planned contrasts via paired-samples t-tests revealed that the significant interaction was specifically due to a significant reduction of the Successful-stop vs. Go difference in the rIFC lesion group (*M* = -.005, *SD* = .116) compared to the matched healthy comparison group (*M* = .126, *SD* = .148), (*t*(15) = -3.862, *p* = .002, *d* = -.965), with a large effect size. The Failed-stop vs. Go difference were also numerically reduced in the rIFC lesion group (*M* = -.018, *SD* = .106) compared to matched healthy comparisons (*M* = .054, *SD* = .150), though that difference was not significant (*t*(15) = -1.720, *p* = .106, *d* = -.430).

The same 2x3 ANOVA for the non-rIFC lesion group and their healthy matched comparisons only revealed the main effect of TRIAL TYPE (*F*(2, 60) = 6.468, *p* = .003, *η*^2^ = .177), with a comparable effect size to the rIFC group. However, unlike for the rIFC group, the interaction between GROUP and TRIAL TYPE and the main effect of GROUP were not significant (Interaction: *F*(2, 60) = .822, *p* = .445, *η*^2^ = .027; GROUP: *F*(1, 30) = .786, *p* = .382, *p* = .026). Furthermore, the same planned contrasts for the Successful-stop vs. Go difference revealed no difference in β-burst rate between non-rIFC lesion group (*M* = .092, *SD* = .092) and matched healthy comparison (*M* = .064, *SD* = .125), (*t*(15) = 1.02, *p* = .324, *d*= .255). Finally, there was no significant difference in Failed-Stop vs. Go difference between non-rIFC lesion group (*M* = .074, *SD* = .122) and their healthy comparisons (*M* = .018, *SD* = .157), (*t*(15) = 1.497, *p* = .155, *d* = .374). Together, these results show that rIFC lesion patients show a significant reduction of stop-related frontal β-bursts compared to healthy comparisons, which was not the case for non- rIFC lesion patients.

We then also directly compared the Stop vs. Go differences in frontal β between the two lesion groups. This comparison again showed a significantly reduced β-burst rate difference between Successful-stop and Go trials in rIFC lesion group (*M* = -.005, *SD* = .116) compared to the non-rIFC lesion group (*M* = .092, *SD* = .092) (*t*(30) = -2.608, *p* = .014, *d* = -.922, *independent samples t-test*). The failed-stop vs. Go difference showed the same pattern, with the rIFC lesion group (*M* = -.018, *SD* = .106) showing a significantly reduced β-burst rate compared to the non- rIFC lesion group (*M* = .074, *SD* = .122), (*t*(30) = -2.265, *p* = .031, *d* = -.801, *independent samples t-test*).

To sum up, these results confirmed our hypothesis that rIFC lesion patients show reduced β-burst rates on successful stop-trials compared to healthy comparisons, given their increase in stop-signal trigger failures. Moreover, this impairment was specific to the rIFC lesion group, as no group difference was found between the non-rIFC lesion group and matched comparisons. Finally, the rIFC group also showed a significant reduction in frontal burst rates when directly compared to the non-rIFC lesion group.

### EEG results: Sensorimotor β-bursts after stop-related frontal bursts

We then investigated the temporal development of sensorimotor β-bursts after frontal β- bursts that occurred during the critical post-stop-signal window on successful-stop trials (***Figure 4***). First, we replicated that across the whole sample, sensorimotor β bursts were increased following frontal β bursts (compared to trials without frontal β bursts). This was done by comparing the sensorimotor β burst rates on successful stop trials that contained frontal β bursts in the stop-signal-to-SSRT period to that of trials without frontal β bursts (in which sensorimotor β bursts were time-locked to a random time point in the stop-signal-to-SSRT period). This analysis showed that sensorimotor β burst rates were significantly increased following frontal β bursts (***Figure 4A***), replicating Wessel (2020). To test for potential group differences in this pattern, we then compared the sensorimotor burst rate between the respective lesion groups and their healthy comparisons (***Figure 4B***). No time point showed any significant differences at p > .05, even absent any corrections for multiple comparisons. Together, this shows that the pattern of increased sensorimotor β-bursts following stop-related frontal β burst was intact, even in the rIFC group.

**Figure 4.**
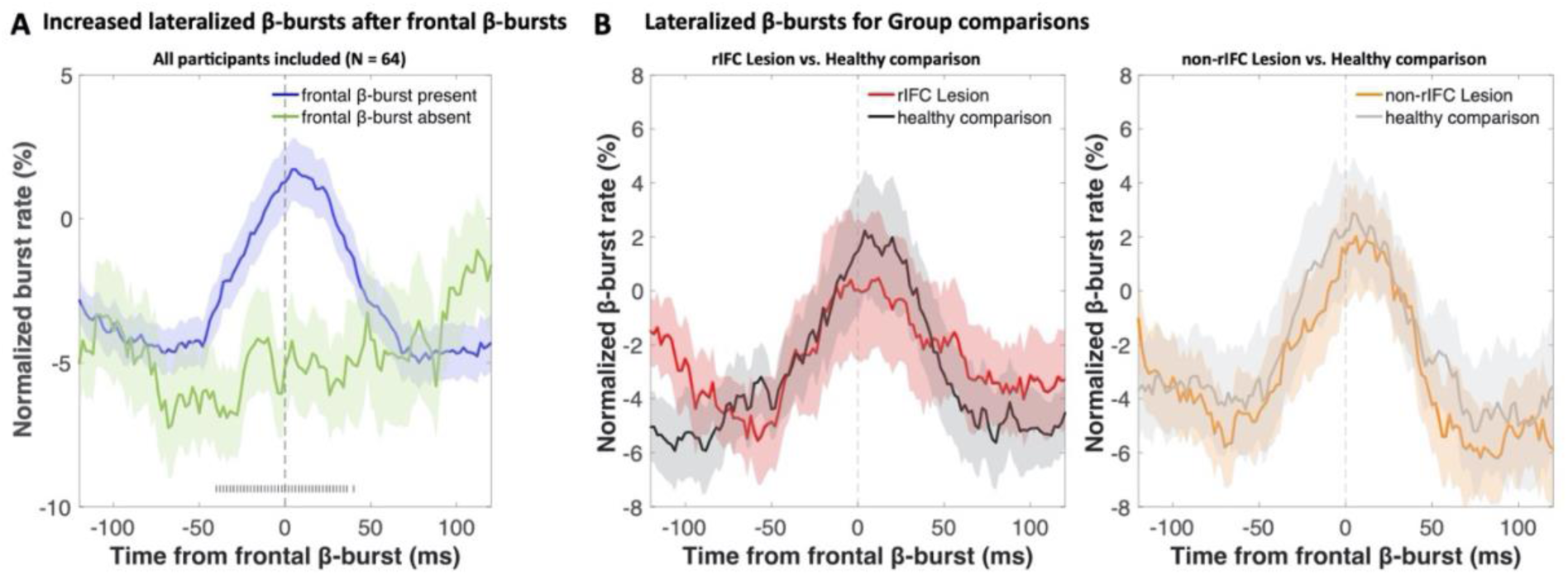
Normalized β-burst rate (%) at bilateral sensorimotor electrodes (C3/C4) following the first frontal β-burst in individual successful stop trials during the critical time period (stop-signal to SSRT). **A**, a comparison between trials with frontal β-bursts to trials without frontal β-bursts (where the sensorimotor β-bursts were time-locked to a random time point within a critical time period). The sensorimotor burst rates following frontal β-bursts were significantly increased (significance outlined at the bottom of the graph). **B**, Group differences (lesion vs. matched healthy comparison) in sensorimotor β-burst rates. No significantly differences were found. The colored patch shows the ±SEM at each timepoint.

## Discussion

In the current study, we re-examined the role that rIFC plays in action-stopping and inhibitory control. rIFC has been at the center of arguably the most influential neural theory of inhibitory control. The proposal that “response inhibition can be localized to a discrete region within of the PFC” (Aron et al., 2003) has been highly influential (Aron et al. 2004). In its most recent iteration, the theory states that rIFC “implements a brake over response tendencies” (Aron et al., 2014). Our work shows that this theory needs further revision. Indeed, rIFC does not seem to be primarily responsible for the *implementation* of inhibitory control. Instead, its role appears to be the triggering or *initiation* of the cascade that ultimately leads to the stopping of action. Concomitantly, rIFC patients showed more than five-fold increase in stop-signal trigger failures, as well as a significant reduction in stop-related β-bursts over frontal cortex. However, the implementation of inhibitory control itself appears to take place in other regions – and appears to remain intact in the rIFC group. In the current study, this was suggested by the fact that in all groups (crucially, including rIFC patients), sensorimotor β-bursts were upregulated when a β-burst in frontal cortex did occur (i.e., when an initiation signal from frontal cortex was successfully sent).

There are at least two possible explanations for the finding that lesions to the rIFC cause such a substantial increase in stop-signal trigger failures. Both explanations imply substantially different roles for rIFC in cognitive control and cognition more broadly. The first possible explanation is that the function of rIFC is to specifically trigger inhibitory control during action-stopping or braking (e.g., in the types of situations that are simulated in the stop-signal task). In this framework, rIFC would still have an inhibition-specific role – though not in the actual implementation of the underlying process, but merely in its initiation. A different, competing explanation is that the function of rIFC is to detect *any* salient signal, and that it fulfills no role that is specific to inhibitory control. In the stop-signal task, rIFC would therefore merely be activated because of the saliency of the stop-signal, but not specifically because of the associated inhibitory requirements. In other words, it is up for debate whether the rIFC fulfills a role that is specific to inhibitory control or stopping/braking, or one that is more domain-general.

Along the lines of the latter explanation, rIFC is prominently at the core of another influential theory, which proposes that rIFC is part of a ventral attention network that functions as a ‘circuit breaker’. In this framework, rIFC’s role is to orient attention towards salient events – which would include, but not be limited to – stop-signals (Corbetta et al., 2008; Corbetta & Shulman, 2002). In line with this theory, fMRI work has shown that similar regions of rIFC are activated not just after stop-signals, but after any sort of salient, infrequent event (Erika-Florence et al., 2014; Sharp et al., 2010). As such, it is possible that the role of rIFC in stop-signal performance is indeed not specific to inhibitory control situations, but instead, that it merely detects the saliency of the stop-signal. In that scenario, the increase in trigger failures in the rIFC lesion group would reflect a more general attentional deficit, which – in the specific case of the stop-signal task – would express itself in the impaired detection of the stop-signal. However, while this impaired detection of the stop-signal would increase SSRT estimates, the underlying cause of that increase would be unrelated to the implementation of inhibitory control.

Crucially, however, any attempt to disentangle the domain-general detection of a salient stimulus from the stopping-specific implementation of inhibitory control is complicated by another factor. Namely, all salient stimuli, even those presented outside of stop-signal contexts, lead to an automatic, physiological inhibition of the motor system (Dutra et al., 2018; Tatz et al., 2021; Wessel, 2018; Wessel & Aron, 2017). For example, salient stimuli lead to a non-selective inhibition of cortico-motor excitability (Iacullo et al., 2020), activate basal ganglia regions involved in inhibitory pathways (Wessel et al., 2016), reduce isometrically exerted force (Novembre et al., 2018, ; 2019), and increase motoric response times (e.g., Parmentier, 2008). Indeed, it appears as if inhibitory control is a ubiquitous part of an organism’s orienting response to salient stimuli (Sokolov, 1963). As such, inhibition and attention may be inextricably linked. If that is indeed the case, both competing theories of rIFC may be partially correct: the role of rIFC in cognitive control may indeed be domain-general, but its domain-general role may be to *trigger inhibitory control following any type of salient stimulus*, specifically as part of a stereotypic and ubiquitous orienting response (Wessel & Aron, 2017). Either way, while it still remains to be seen whether the rIFC does have a specific role in inhibitory control or not, it seems safe to conclude that its role is not in the implementation of that process.

This latter fact is evident in the current study by the fact rIFC patients could appropriately implement inhibitory control on the subset of trials in which it was ostensibly triggered successfully. This is primarily supported by the β-burst patterns. β activity in the LFP has long been linked to movement regulation (e.g., Kilavik et al., 2013; Swann et al., 2009; 2011). Recent work has shown that β activity in the human (and non-human) local field potential occurs in clearly demarcated, transient bursts (Feingold et al., 2015; Sherman et al., 2016; Shin et al., 2017). In acute stopping situations, a short-latency increase in such β-bursts over frontal cortex takes place (cf., ***Figure 3***, see also: Enz et al., 2021; Jana et al., 2020; Wessel, 2020). These frontal bursts are then following by a near-immediate increase in β-burst rates over sensorimotor cortex (cf., ***Figure 4***, see also: Diesburg et al., 2021; Wessel, 2020). Sensorimotor β has been long proposed to reflect an inhibited state of the motor system at rest (Kilavik et al., 2013, Soh et al., 2021). A re-instantiation of this sensorimotor β-bursting after frontal β-bursts has therefore been proposed to reflect the final stage of the inhibitory cascade during rapid action-stopping – the return of the motor system to its inhibited default state (Wessel, 2020). In the current study, we found that the initial frontal β-bursts immediately after the stop-signal were substantially reduced in the rIFC lesion group – both compared to healthy comparisons and compared to non-rIFC lesion patients. Conversely, however, all groups – notably including rIFC lesion patients – showed a significant increase in sensorimotor β-bursting immediately after frontal β-bursts. In other words, in cases in which a frontal β-burst did take place in rIFC patients, their motor system was successfully return to its inhibited state.

Together, this suggests that inhibitory control, while perhaps triggered in rIFC, is implemented in other areas. While there has been vigorous debate about the role of the rIFC (and other cortical regions, such as the pre-SMA, (Nachev et al., 2007) in action-stopping and inhibitory control, there seems to be some degree of consensus that the final steps on the way to successful action-stopping involve an interruption of thalamocortical motor representations via the output nuclei of the basal ganglia (see Jahanshahi et al., 2015 for a review). This interruption is likely caused by the subthalamic nucleus (which is part of two long-hypothesized anti-kinetic cortico-basal ganglia-thalamocortical pathways: indirect and hyper-direct, cf., (Nambu et al., 2002; Parent & Hazrati, 1995). In line with this, recent work has tracked the abovementioned β-burst dynamics further along the basal ganglia regions that are purportedly involved in the implementation of inhibitory control. Indeed, similar to the frontal β-burst rates report here and elsewhere, β-burst rates are also increased after stop-signals in both the subthalamic nucleus and the motor regions of the thalamus (Diesburg et al., 2021). Moreover, subthalamic bursts in particular are also followed by short-latency upregulations of β-bursts in sensorimotor areas, just like the frontal bursts in the current study (and others). As such, it seems likely that while the rIFC is not directly involved in implementing inhibitory control, that function is fulfilled by the basal ganglia regions of the hyper and/or indirect pathways (Schmidt et al., 2013). To this point, recent work using electrical stimulation has shown that there is a direct, monosynaptic connection between rIFC and the subthalamic nucleus, and that this connection is highly relevant for the speed of inhibitory control (Chen et al., 2020). Together, a coherent picture emerges according to which rIFC, once activated by a salient signal such as the stop-signal, triggers a multi-step cascade that culminates in successful action-stopping. At a minimum, this cascade involves the subthalamic nucleus, the output nuclei of the basal ganglia, and the motor thalamus, with the implementation of inhibitory control taking place in this subcortical chain. As these subcortical areas were intact in our rIFC lesion patients, it makes sense for the implementation of inhibitory control – once triggered – to be unimpaired in this population.

Taken together, our data show that the dominant model of inhibitory control in the human brain needs to be fundamentally revised. Rather than the rIFC “implementing a brake over response tendencies”, it appears as though its primary role is to detect salient signals (such as stop-signals) and then trigger the inhibitory control process, with implementation taking place in other areas.

## Materials and Methods

### Participants

Sixty-four participants across four groups of N = 16 were recruited for the study, matching the sample size of the rIFC lesion group in the original Aron et al. (2003) investigation. In addition to patients with focal lesions within the rIFC, we also included 16 non-rIFC lesion comparison patients – i.e., individuals with lesions outside of rIFC (see ***Figure 1*** for lesion overlap maps). All rIFC lesion and non-rIFC lesion patients were selected from the Neurological Patient Registry of the University of Iowa’s Division of Behavioral Neurology and Cognitive Neuroscience. Thirty-two age- and sex-matched healthy comparison participants were then also recruited from the Cognitive Neuroscience Registry for Normative Data of the Division of Behavioral Neurology and Cognitive Neuroscience and through local ads. All participants received detailed information describing the experiment and provided informed consent prior to participating in the study. The study was approved by the Institutional Review Board at the University of Iowa and conducted in accordance with the Declaration of Helsinki. Data collection was performed in an EEG laboratory in the Neurology Clinic at the University of Iowa between October 2018 and July 2021. All lesion patients were compensated at an hourly rate of $30, while healthy participants were compensated at an hourly rate of $15. Monetary compensations for mileage and meal were also provided. Demographic data for all participants are presented in Table 1.

### Task and procedure

Participants performed a stop-signal task presented using Psychtoolbox (version 3, Brainard, 1997) in MATLAB 2015b (TheMathWorks, Natick, MA) on an Ubuntu Linux desktop computer. Responses were made using a standard QWERTY USB keyboard. A schematic design of the stop-signal task is shown in ***Figure 2***. Each trial began with a fixation cross at the center of the screen (500 ms) followed by a black arrow (Go-signal) pointing either left or right, displayed for 1000 ms. Two white key stickers were attached on the “q” key and “p” key in the keyboard to indicate Go stimulus-response key mapping. Participants were instructed to press the left key (the “q” key) with their left index finger in response to the left arrow and the right key (the “p” key) with their right index finger in response to the right arrow as fast and accurately as possible. In five rIFC lesion patients (as well as their respective matched comparison participants) participants instead made unimanual responses with two fingers of their right hand using the arrow keys on the keyboard. This was due to lesions in their right hemisphere that encompassed motor cortex and surrounding areas, which could affect the mobility of their left hand.

If no response was made during the Go-signal presentation (1000 ms), the feedback “Too Slow!” message was presented for 1000ms at the center of the screen. Note that no response times were collected after the end of this 1000 ms window for rIFC patients and their healthy comparisons, as well as for four non-rIFC lesion patients and two healthy comparisons (the procedure was then slightly changed to keep recording responses even after this window, though those responses were still counted as misses for the purposes of the behavioral analyses). While our ex-Gaussian approach (see below) was explicitly designed to handle the potential censoring of the Go-RT distributions resulting from this cutoff deadline, we still re-collected two rIFC lesion patients’ data, whose Go-RT distributions showed signs of censoring.

On one-third of trials, an auditory Stop-signal (900Hz sine-wave tones of 100 ms duration) was presented after the Go-signal at a varying stop-signal delay (SSD). Participants were instructed to withhold their response on such trials. The SSD was initially set to 200 ms and adjusted separately for left and right responses depending on stop success (50 ms increment) or failure (50 ms decrement), with a goal of achieving an overall p(stop) of approximately .50 (Verbruggen et al., 2019). The overall trial length was fixed to 3,500 ms. Before the experiment, participants performed a short practice block (24 trials). In total, participants underwent 480 trials (eight blocks of 60 trials; 320 go / 160 stop-trials overall). In each rest period between blocks, performance feedback was given on the previous block.

### Bayesian modeling of behavioral data

The stop-signal data were analyzed with BEESTS, a hierarchical Bayesian modeling technique that simultaneously accounts for the shape of Go-RT and SSRT distributions and the prevalence of trigger failures in the stop-signal task (Matzke et al., 2013; Matzke, Love, et al., 2017). As shown in ***Figure 2B***, BEESTS is based on the horse-race model (Logan & Cowan, 1984) and assumes that response inhibition depends on the relative finishing times of a go and a stop runner, triggered by the go and the stop stimuli, respectively. On a given trial, if the go runner is slower than SSD + the finishing time of the stop runner, the go response is successfully stopped (i.e., stop process wins the race). If the go runner is faster than SSD + the finishing time of the stop runner, inhibition fails and a signal-respond RT (e.g., grey-color distribution in ***Figure 2***) is produced. BEESTS assumes that the finishing times of the go (Go-RT distribution) and the stop runners (SSRT distribution) follow an ex-Gaussian distribution with parameters ***µ***, ***σ***, and ***τ***. The ***µ*** and ***σ*** parameters reflect the mean and the standard deviation of the Gaussian component and ***τ*** gives the mean of the exponential component and reflects the slow tail of the distribution. The mean and variance of the finishing time distributions can be obtained as ***µ*** + ***τ*** (i.e., mean Go-RT and SSRT) and ***σ*** ^2^ + ***τ*** ^2^, respectively. Using a mixture-likelihood approach, the model can be augmented with a parameter, P(*TF*), that quantifies the probability that participants fail to trigger the stop runner (Matzke, Love, et al., 2017).

Incorrect RTs and RTs faster than 200 ms (e.g., anticipatory responses) were removed before fitting the data. In total, this excluded 111 trials across the rIFC group (1.45% of all data), 15 trials in the non-rIFC group (.2% of all data) and 88 trials in the matched comparisons (.55% of all data). We treated omissions on go trials and RTs on go and stop-signal trials slower than 1000 ms as censored observations and adjusted the likelihood of the model to account for the upper censoring of the finishing time distributions. We modeled the data of each group separately using a single go runner and a stop runner, resulting in seven parameters per participant: ***µgo***, ***σgo***, and ***τgo*** for the go runner, and ***µstop***, ***σstop***, ***τstop***, and P(TF) for the stop runner. The P(TF) parameter was estimated on the real line after transformation from the probability scale using a probit transformation. We did not use the recently extended version of the model (which uses separate go runners corresponding to the two go response options) as direction error responses were very rare (Matzke et al., 2019). Note also that the “censored” BEESTS model, in its current form, does not allow for the estimation of “go failures”, i.e., the probability that participants fail to trigger the go runner.

We assumed (truncated) normal population-level distributions for all model parameters, including the probit transformed P(TF) parameter. The population-level distributions were parametrized and estimated in terms of their location and scale parameters, which were then transformed back to means and standard deviations for inference. As shown in the Supplementary Materials, we assigned weakly informative priors to the population-level location and scale parameters that covered a wide but realistic range (e.g., Matzke et al., 2019).

The analyses were carried out in the Dynamic Models of Choice software (Heathcote et al., 2019) using Differential Evolution Markov chain Monte Carlo (DE-MCMC; Ter Braak, 2006) sampling implemented in the R programming environment (R Core Team, 2015). We used parameter estimates obtained from fitting each participant’s data individually using non-hierarchical Bayesian estimation as start values for the hierarchical sampling routine. The number of MCMC chains was set to 21, i.e., three times the number of participant-level model parameters. To reduce autocorrelation, the MCMC chains were thinned to retain only every 15^th^ draw from the joint posterior distribution. During the burn-in period, the probability of a migration step was set to 5%, after which only crossover steps were performed. Convergence was assessed using univariate and multivariate proportional scale-reduction factors (*R̂* < 1.1; Brooks & Gelman, 1998; Gelman & Rubin, 1992) and visual inspection of the MCMC chains.

The absolute goodness-of-fit of the model was assessed with posterior predictive model checks (Gelman et al., 1996) using the average cumulative distribution function of Go-RTs and signal-respond RTs, inhibition functions, and median signal-respond RTs as a function of SSD. Decision about the descriptive accuracy of the model was based on visual inspection of the model predictions, aided with posterior predictive *p* values (for details, see Matzke et al., 2019). As shown in the Supplementary Materials, the model provided a good account of all these aspects of the observed data for the rIFC lesion patients and the matched healthy comparison participants (***Figures S1-S4***), but it showed a quantitatively small misfit to the average inhibition function at short SSDs in the other two groups (***Figures S6*** and ***S8***). To examine the robustness of the results to a possible model misspecification, we sequentially removed all stop-signal trials from the data of the non-rIFC lesion and the matched comparison group at SSDs of 0, 50, 100, 150, and 250 ms, re-fit the model, and re-assessed the model’s descriptive accuracy. Descriptive accuracy improved as stop-signal trials on short SSDs were removed (***Figure S10*** and ***12***). To avoid overconfidence and to ensure that our conclusions are based on a descriptively accurate model, we report results based on mixing the posterior distributions estimated using the full data set and the five subsets after removing stop-signal delays between 0 and 250 ms.

### Statistical inference on model parameters

We used the mean of the posterior distributions as point estimate for the BEESTS model parameters, and the 2.5^th^ and 97.5^th^ percentile of the distributions (i.e., 95% credible interval, CI) to quantify estimation uncertainty. Inference about group differences was based on the overlap between the posterior distributions of the population-level parameters of the different groups (i.e., both lesion groups vs. their respective matched healthy comparison group). Overlap was quantified using Bayesian *p* values computed as the proportion of samples in the posterior distribution of the lesion group that is larger than in the matched comparison group. Bayesian *p* values close to 0 or 1 indicate that the posterior distribution of the lesion group is shifted to lower or higher values, respectively, relative to the matched comparison group, suggesting the presence of a group difference. Bayesian *p* values were computed after appropriate transformations of the posterior distributions (i.e., bivariate inverse probit transformation for the P(TF) parameter and transformation of the truncated normal population-level location parameter to means). The posterior distributions of the population-level mean Go-RT and SSRT were obtained by computing ***µgo*** + ***τgo*** and ***µstop*** + ***τstop,*** respectively, for each MCMC iteration and then collapsing the resulting population-level samples in a single distribution across chains. Similarly, the posterior distributions of the participant-level mean Go-RT and SSRT were computed by summing the corresponding participant-level ***µ*** and ***τ*** samples for each iteration and collapsing the resulting samples across the chains. The point estimates used in the analysis of β-burst events reflect the mean of the participant-level posteriors.

The preregistration document for these analyses can be found here: https://osf.io/d9r4s/.

### EEG Recording

64-channel EEG data in the extended 10-10 system were recorded using two Brain Products systems (actiChamp or MRplus). Ground and reference electrodes were placed at AFz and Pz, respectively. The MRplus system included two additional electrodes on the left canthus (for the horizontal eye movement) and below the left eye (for the vertical eye movement). EEG data was digitized with a sampling rate of 500 Hz, with hardware filters set to 10 seconds time-constant high-pass and 1000 Hz low-pass.

### EEG preprocessing

EEG data preprocessing was conducted using custom routines in MATLAB. The data were filtered (high pass cut-off: .3Hz; low pass-cutoff: 50Hz) and then visually inspected to identify and remove non-stereotypical artifacts. The data were subsequently re-referenced to common average and subjected to a temporal infomax independent component analysis (ICA; Bell & Sejnowski, 1995) as implemented in EEGLAB (Delorme & Makeig, 2004). Components representing stereotyped artifact activity (saccades, blinks, and electrode artifacts) were identified using outlier-based statistics and were removed from the data. We further obtained dipole solutions for ICs using the Dipfit plug-in for EEGLAB and further rejected ICs with residual variance larger than 15%, which typically represent non-brain data (Delorme et al., 2012). The remaining components were backprojected into channel space to reconstruct artifact-free channel data and subjected to further analyses. Finally, the channel-space data were then transformed using the current-source density method (CSD; Kayser & Tenke, 2006), which attenuates the effects of volume conduction on the scalp-measured activity.

### β-burst detection

β-burst detection was performed using the same method as described in Wessel (2020), except with a burst detection threshold of 2X median power (rather than 6x median), following the recent recommendation from Enz et al. (2021). First, each electrode’s data were convolved with a complex Morlet wavelet of the form:

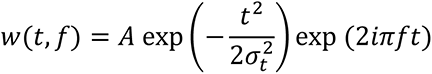

With 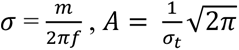, and m = 7 (cycles) for each of 15 evenly spaced frequencies spanning the beta-band (15-29 Hz). Time-frequency power estimates were extracted by calculating the squared magnitude of the complex wavelet-convolved data. These power estimates were then epoched relative to the events in question (ranging from −500 to +1000 ms with respect to Stop-/Go-signals). Individual β-bursts were defined as local maxima (using the MATLAB function *imregionalmax*) in the trial-by-trial β-band time-frequency power matrix for which the power exceeded a cutoff of 2X the median power of the entire time-frequency power matrix for that electrode (Enz et al., 2021).

### Statistical analysis of β-burst events

The quantification of frontal β-bursts was done as in Wessel (2020). β-burst events in the critical time period ranging from the stop-signal onset to each individual’s SSRT estimate were counted separately for successful- and failed-Stop trials at electrode FCz. For Go trials, we counted the number of β-burst events in a time period of identical length, ranging from the current stop-signal delay on the trial and the participants’ SSRT estimate (in other words, in the time period during which a stop-signal would have appeared on that particular trial and the end of SSRT). We then converted these numbers to β-burst rate (bursts per second) by dividing each participant’s burst rates with the length of individual SSRT estimate. We then normalized each participant’s burst/sec measurement with a baseline time period of [-500 0] relative to the Go stimulus onset on the trial for each trial-type (Successful Stop, Failed Stop and Go). These normalized β-burst rates were then analyzed with 2-by-3 mixed ANOVA with between-subjects factor of LESION [lesion vs. matched healthy comparison] and within-subjects factor of TRIAL TYPE [Successful stop, Failed stop, Go]. This was done separately for the rIFC lesion group and the non-rIFC lesion group. Planned comparisons for Stop vs. Go difference (e.g., Successful Stop vs. Go and Failed-Stop vs. Go) within the groups (e.g., lesion vs. matched healthy comparison) were made using paired-samples t-tests.

Furthermore, we investigated the implementation of inhibitory control at sensorimotor sites following frontal β-bursts. To this end, we first identified successful stop-trials in which at least one frontal β-burst event occurred within the critical time period (i.e., between SSD and SSD+SSRT). Next, we time-locked the data to the first frontal β-burst event within that period and quantified the sensorimotor β-burst rate at C3/C4 electrodes from -100 to 100 ms around that frontal burst. In past investigations, this showed clearly increases in β-burst rate immediately (within ∼25 ms) following frontal bursts, which are not present during matched time periods on trials without frontal bursts (e.g., Enz et al., 2021; Wessel, 2020; Diesburg et al., 2021. In the current study, instead of making arbitrary time-bins, we used a sliding window approach, with a search window of 50 ms around each sample point (±25 ms). This method avoids the inherently arbitrary nature of the binning approach.

We first transformed the mean sensorimotor β-burst rates at each of these timepoints to percent change from baseline, with the baseline being the mean β-burst rate in the 500 ms pre-GO signal period of the same trial. Next, these normalized β-burst rates at each time point were compared between groups (lesion vs. matched healthy comparison) using sample-wise paired-samples t-tests covering the whole time period from -100 to 100 ms around the frontal burst.

### Data availability

All data and analysis scripts will be deposited publicly on the OSF upon acceptance of the manuscript.

## Acknowledgements

NA

## Prior Distributions

We modeled the parameters of each participant *j, j=1, …, 16*, in each group using (truncated) normal population-level distributions, with location (M), scale (S), and lower and upper bounds as specified below:

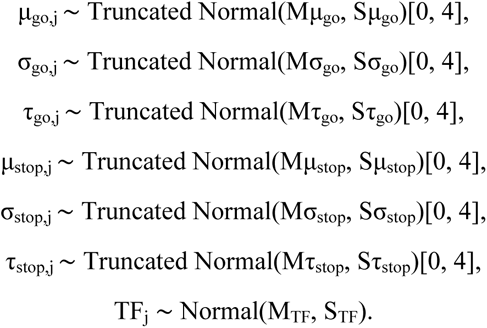

Note that the participant-level P(TF) parameter was first projected from the probability scale to the real line with a probit transformation.

We used (truncated) normal prior distributions for the population-level location parameters:

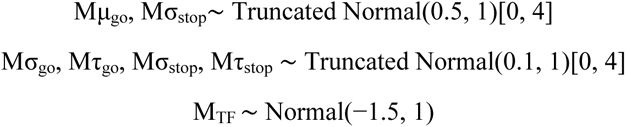

We used exponential prior distributions for the population-level scale parameters:

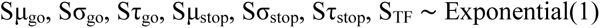

As the data were fit in seconds, the priors are also parametrized on the second scale.

## Goodness-of-Fit

We assessed the descriptive accuracy of the “censored” BEESTS model by comparing the observed data to predictions based on the joint posterior distribution of the participant-level parameters. For each group, we used 500 randomly selected parameter vectors from the participant-level joint posterior to generate 500 predicted stop-signal data sets per participant, using the observed stop-signal delays (SSD) and the observed number of go and stop-signal trials. We focused on three aspects of the data: the average cumulative distribution function of Go-RTs and signal-respond RTs, inhibition functions, and median signal-respond RTs as a function of SSD. The model provided a good account of all these aspects of the data for the rIFC lesion patients and the matched healthy comparison participants (***Figures S1-S4***), but it showed a quantitatively small misfit to the average inhibition function at short SSDs in the other two groups (***Figures S6*** and ***S8***). Descriptive accuracy in the non-rIFC lesion and matched comparison group improved after stop-signal trials on short SSDs were removed (***Figures S10*** and ***S12***).

---

**Figure S1.**
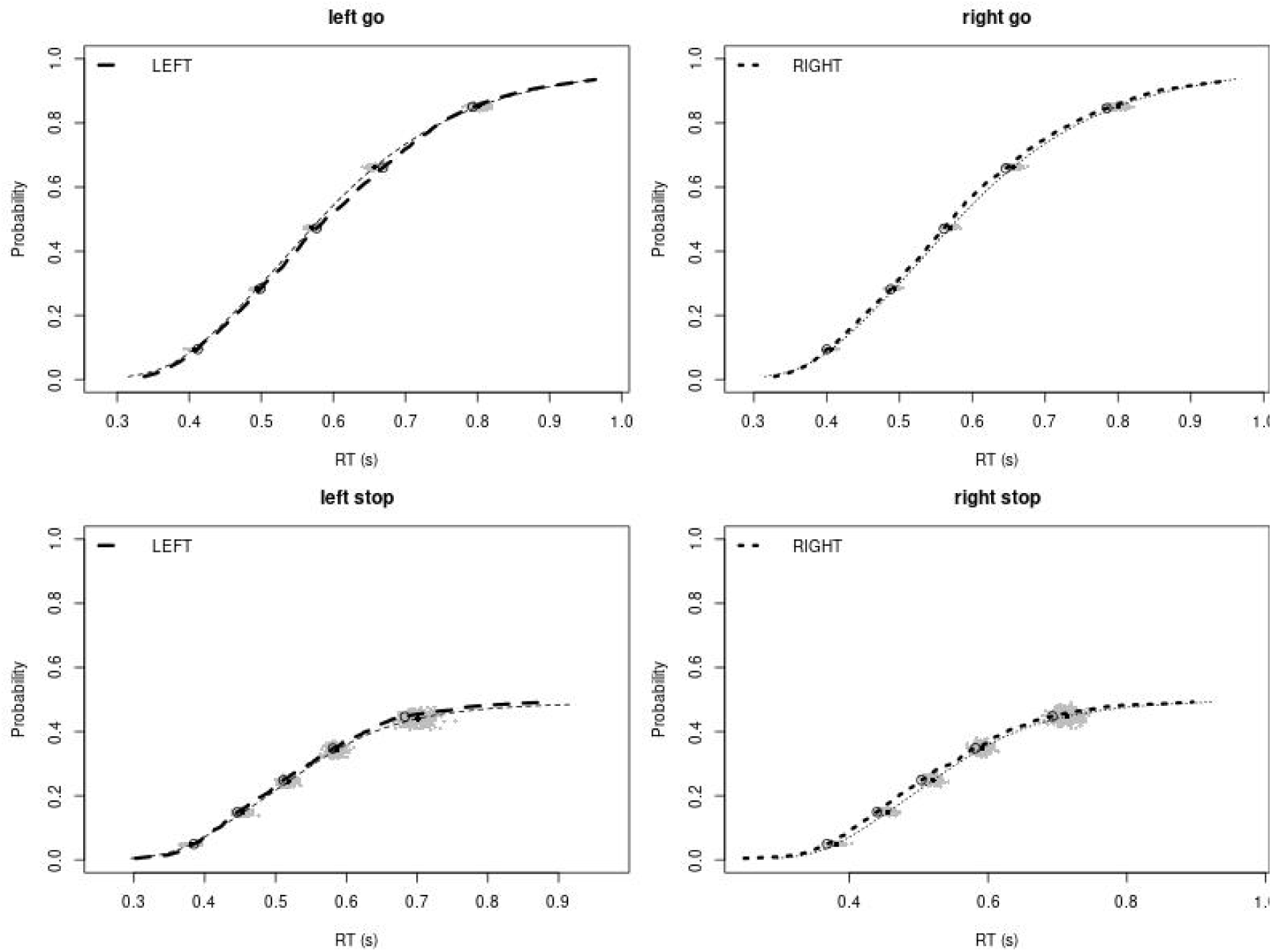
Observed and predicted cumulative distribution function (CDF) of Go-RTs (upper panels) and signal-respond RTs (lower panels), separately for left and right stimuli, for the rIFC lesion group. The observed and predicted CDFs were averaged across participants. Signal-respond RTs were collapsed across SSD. Thick dashed and dotted lines show the CDF of the observed “LEFT” and “RIGHT” responses, respectively. Circles show the 10^th^, 30^th^, 50^th^, 70^th^, and 90^th^ percentile of the distributions. Thin dashed and dotted lines show the CDF of the predicted “LEFT” and “RIGHT” responses, respectively, averaged across the 500 predictions. For each percentile, the gray clouds show the 500 predicted percentiles.

**Figure S2.**
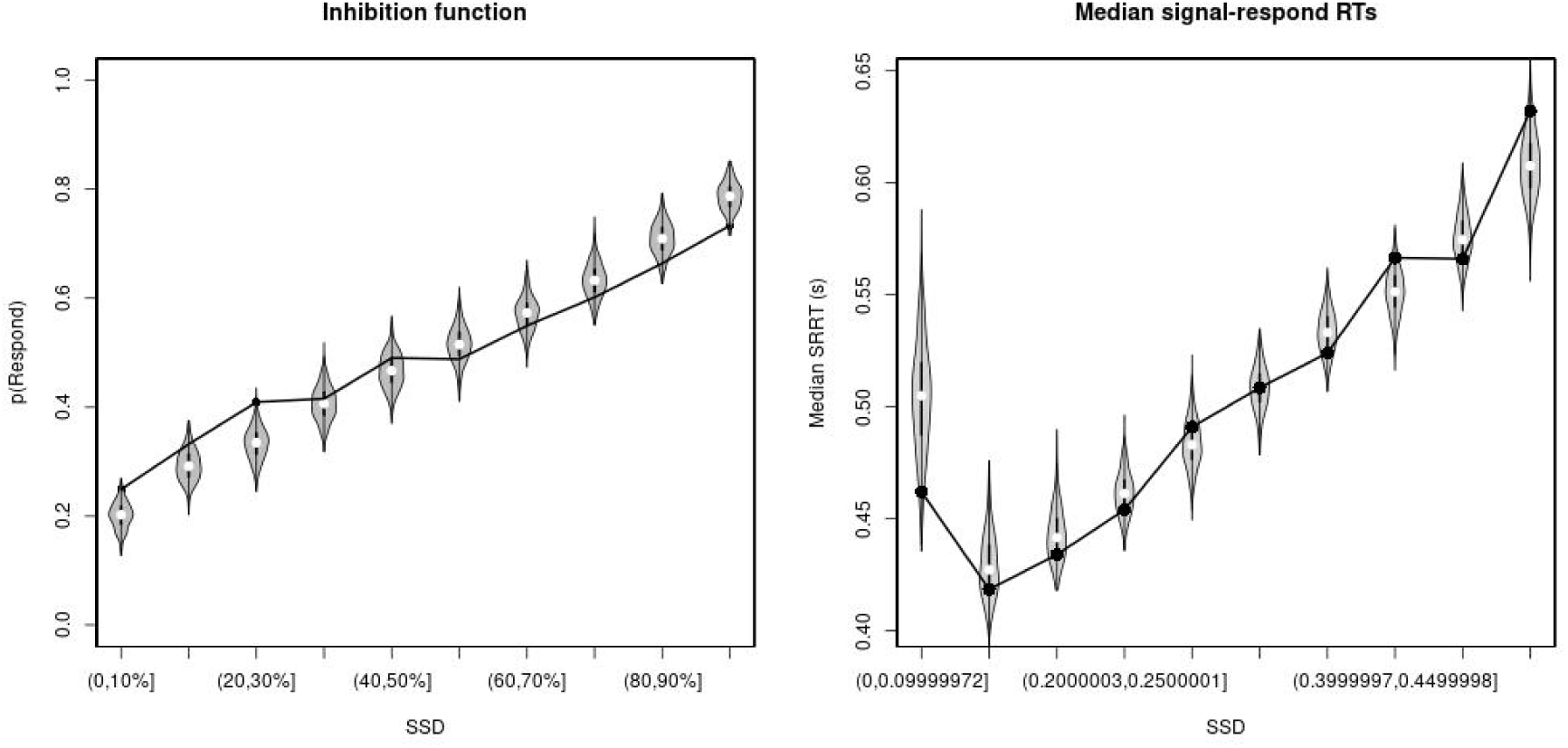
Observed and predicted inhibition function (left panel) and median signal-respond RT as a function of stop-signal delay (SSD; right panel) for the rIFC lesion group. In the left panel, black bullets show the observed average response rate on stop-signal trials (P(Respond)) for each SSD category, where the SSD categories were defined in terms of the percentiles of the distribution of SSDs for each participant and then averaged across participants. In the right panel, black bullets show the observed average median signal-respond RT (SRRT) for each SSD category, where SSD categories were defined by pooling SSDs over participants before calculating the percentiles. The gray violin plots show the distribution of the 500 average response rates and SRRTs predicted by the model, with the white circles representing the median of the predictions.

**Figure S3.**
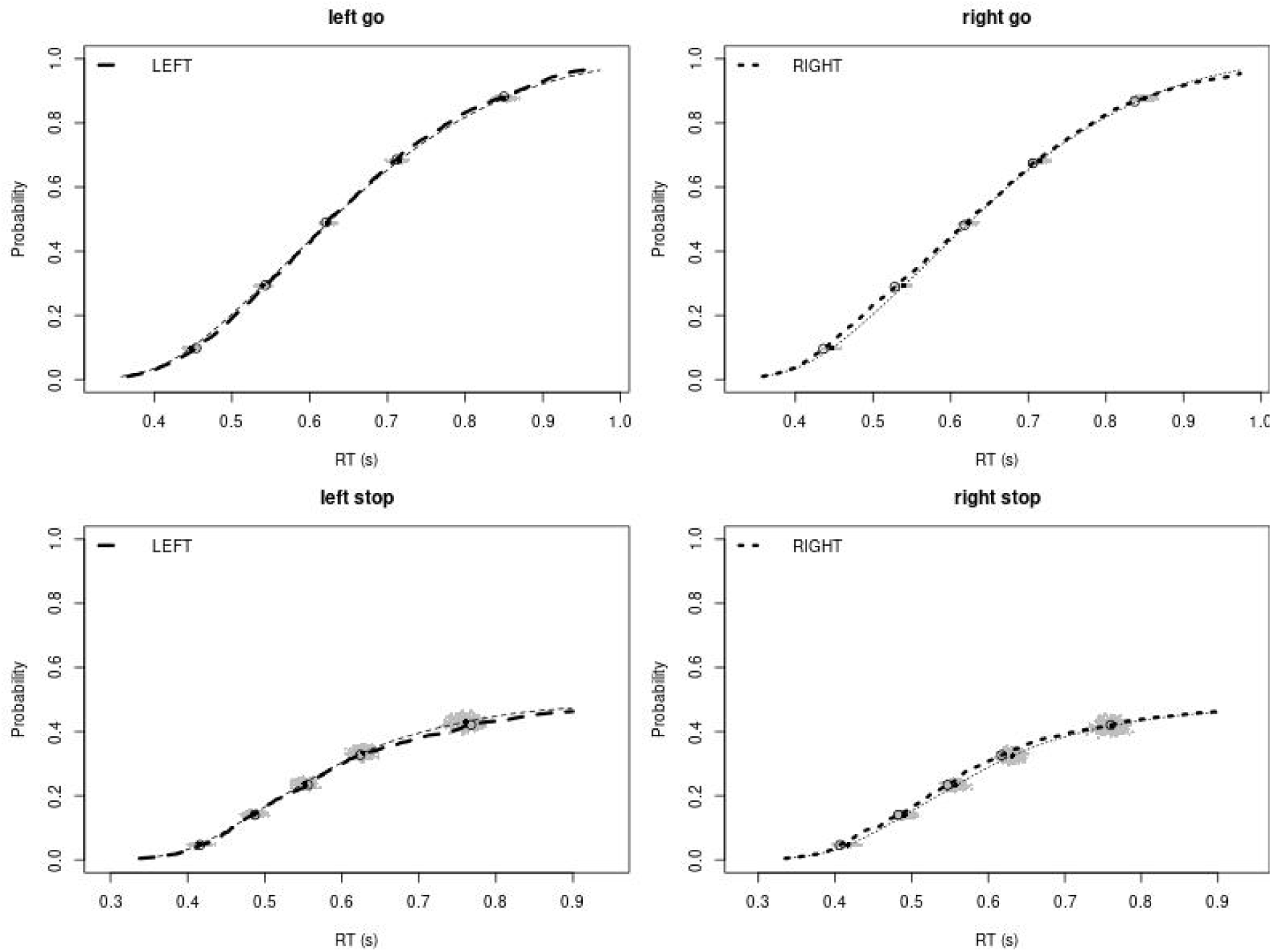
Observed and predicted cumulative distribution function (CDF) of Go-RTs (upper panels) and signal-respond RTs (lower panels), separately for left and right stimuli, for the matched comparison group for the rIFC lesion patients. The observed and predicted CDFs were averaged across participants. Signal-respond RTs were collapsed across SSD. Thick dashed and dotted lines show the CDF of the observed “LEFT” and “RIGHT” responses, respectively. Circles show the 10^th^, 30^th^, 50^th^, 70^th^, and 90^th^ percentile of the distributions. Thin dashed and dotted lines show the CDF of the predicted “LEFT” and “RIGHT” responses, respectively, averaged across the 500 predictions. For each percentile, the gray clouds show the 500 predicted percentiles.

**Figure S4.**
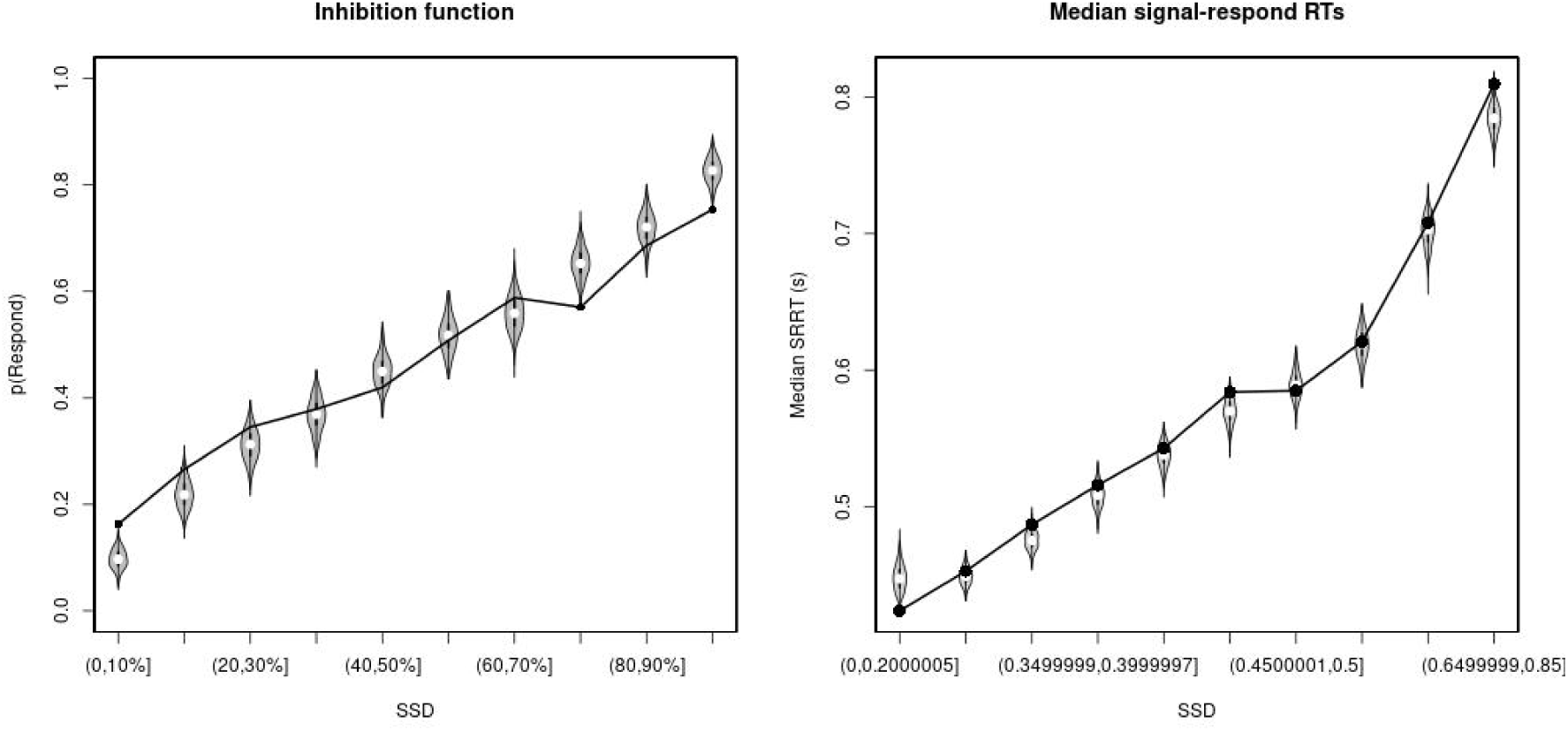
Observed and predicted inhibition function (left panel) and median signal-respond RT as a function of stop-signal delay (SSD; right panel) for the matched comparison group for the rIFC lesion patients. In the left panel, black bullets show the observed average response rate on stop-signal trials (P(Respond)) for each SSD category, where the SSD categories were defined in terms of the percentiles of the distribution of SSDs for each participant and then averaged across participants. In the right panel, black bullets show the observed average median signal-respond RT (SRRT) for each SSD category, where SSD categories were defined by pooling SSDs over participants before calculating the percentiles. The gray violin plots show the distribution of the 500 average response rates and SRRTs predicted by the model, with the white circles representing the median of the predictions.

**Figure S5.**
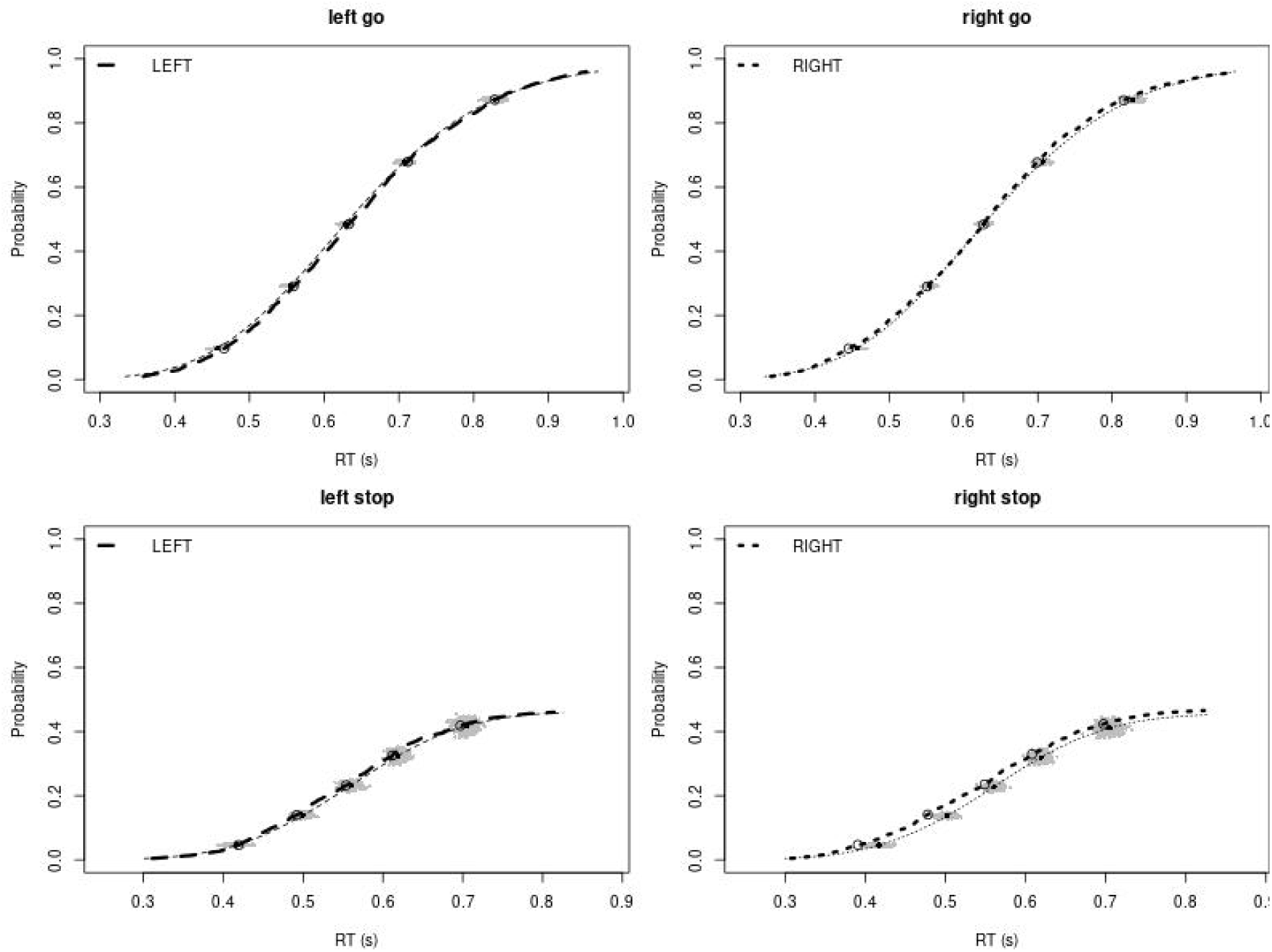
Observed and predicted cumulative distribution function (CDF) of Go-RTs (upper panels) and signal-respond RTs (lower panels), separately for left and right stimuli, for the non-rIFC lesion group. The observed and predicted CDFs were averaged across participants. Signal-respond RTs were collapsed across SSD. Thick dashed and dotted lines show the CDF of the observed “LEFT” and “RIGHT” responses, respectively. Circles show the 10^th^, 30^th^, 50^th^, 70^th^, and 90^th^ percentile of the distributions. Thin dashed and dotted lines show the CDF of the predicted “LEFT” and “RIGHT” responses, respectively, averaged across the 500 predictions. For each percentile, the gray clouds show the 500 predicted percentiles.

**Figure S6.**
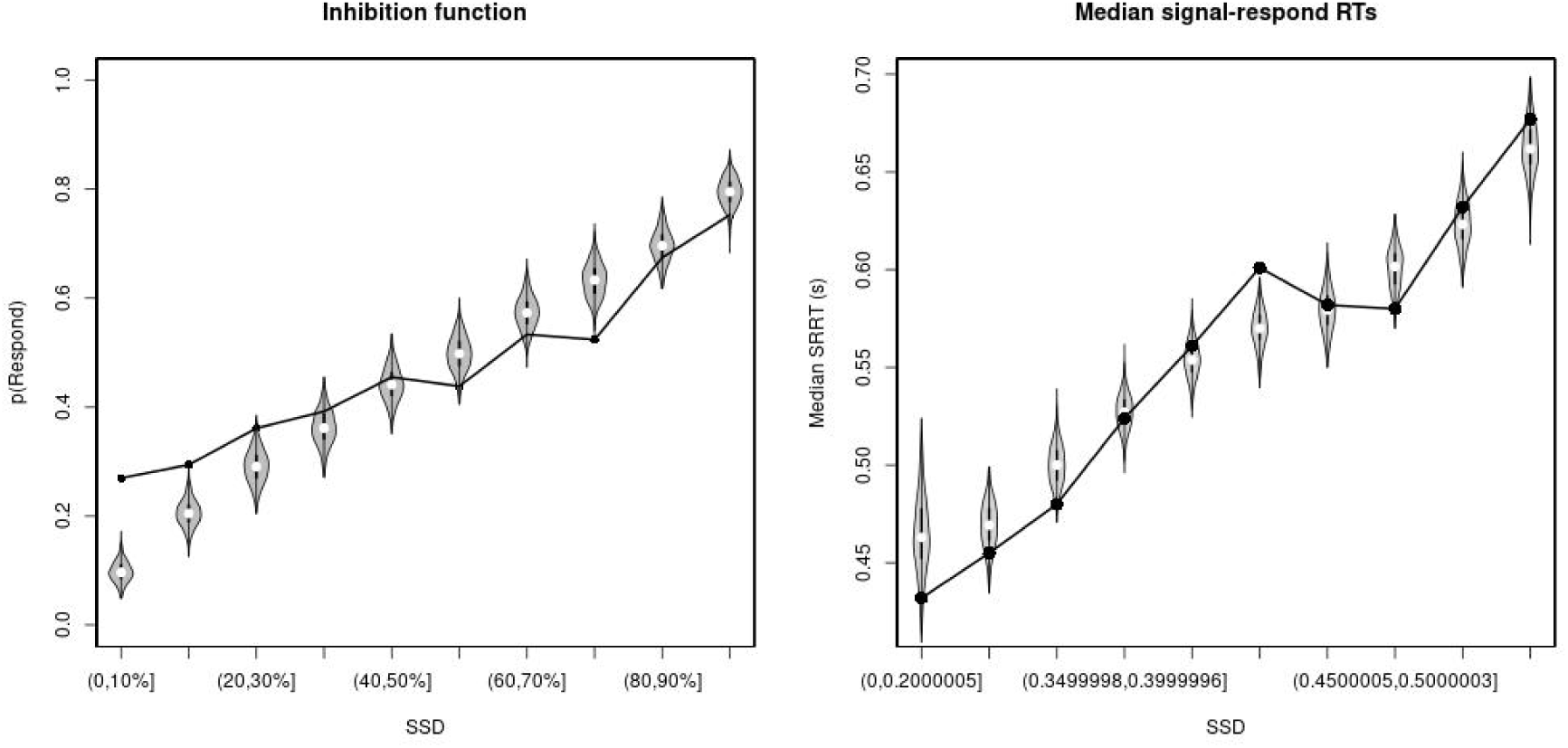
Observed and predicted inhibition function (left panel) and median signal-respond RT as a function of stop-signal delay (SSD; right panel) for the non-rIFC lesion patients. In the left panel, black bullets show the observed average response rate on stop-signal trials (P(Respond)) for each SSD category, where the SSD categories were defined in terms of the percentiles of the distribution of SSDs for each participant and then averaged across participants. In the right panel, black bullets show the observed average median signal-respond RT (SRRT) for each SSD category, where SSD categories were defined by pooling SSDs over participants before calculating the percentiles. The gray violin plots show the distribution of the 500 average response rates and SRRTs predicted by the model, with the white circles representing the median of the predictions.

**Figure S7.**
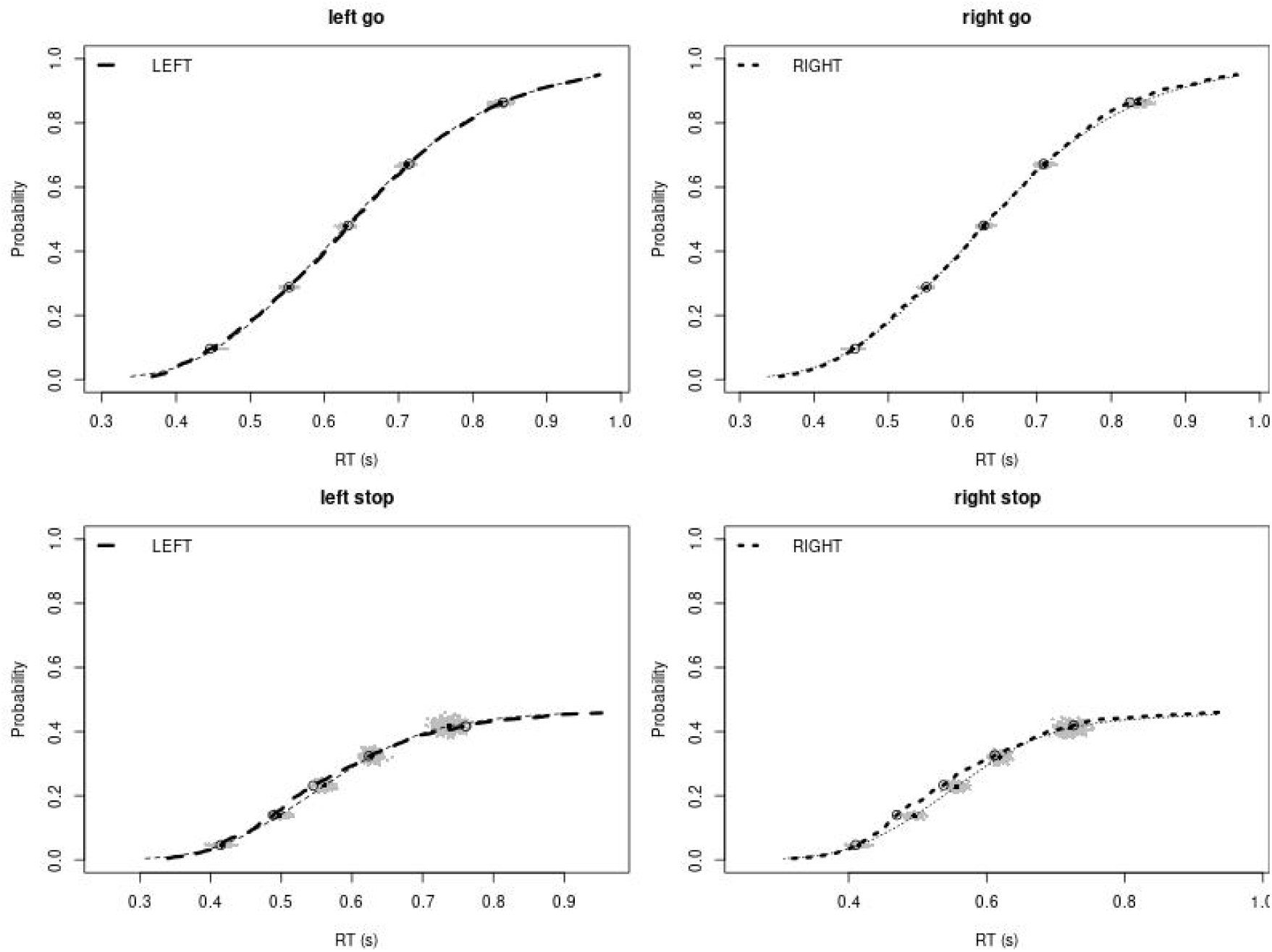
Observed and predicted cumulative distribution function (CDF) of Go-RTs (upper panels) and signal-respond RTs (lower panels), separately for left and right stimuli, for the matched comparison group for the non-rIFC lesion patients. The observed and predicted CDFs were averaged across participants. Signal-respond RTs were collapsed across SSD. Thick dashed and dotted lines show the CDF of the observed “LEFT” and “RIGHT” responses, respectively. Circles show the 10^th^, 30^th^, 50^th^, 70^th^, and 90^th^ percentile of the distributions. Thin dashed and dotted lines show the CDF of the predicted “LEFT” and “RIGHT” responses, respectively, averaged across the 500 predictions. For each percentile, the gray clouds show the 500 predicted percentiles.

**Figure S8.**
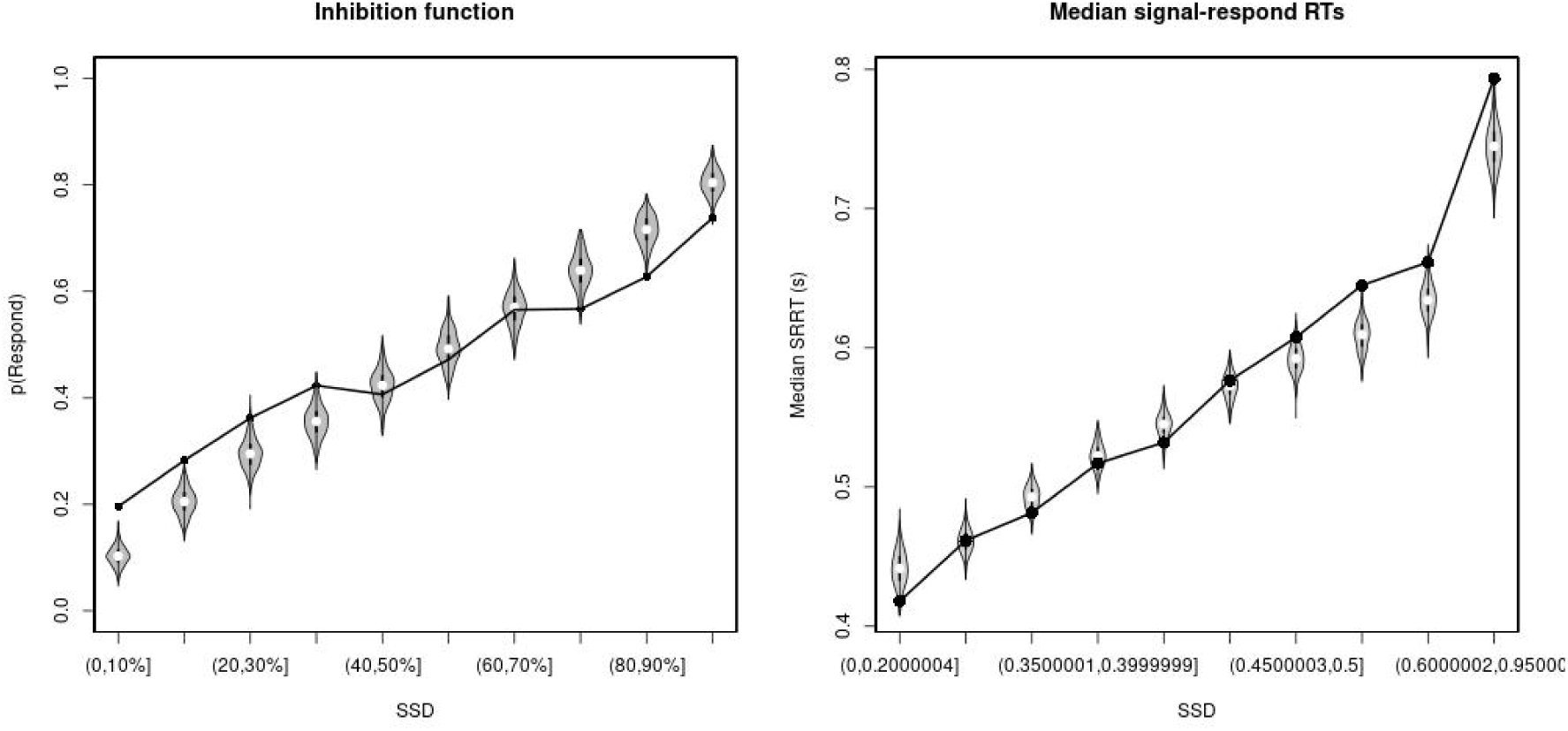
Observed and predicted inhibition function (left panel) and median signal-respond RT as a function of stop-signal delay (SSD; right panel) for the matched comparison group for the non-rIFC lesion patients. In the left panel, black bullets show the observed average response rate on stop-signal trials (P(Respond)) for each SSD category, where the SSD categories were defined in terms of the percentiles of the distribution of SSDs for each participant and then averaged across participants. In the right panel, black bullets show the observed average median signal-respond RT (SRRT) for each SSD category, where SSD categories were defined by pooling SSDs over participants before calculating the percentiles. The gray violin plots show the distribution of the 500 average response rates and SRRTs predicted by the model, with the white circles representing the median of the predictions.

**Figure S9.**
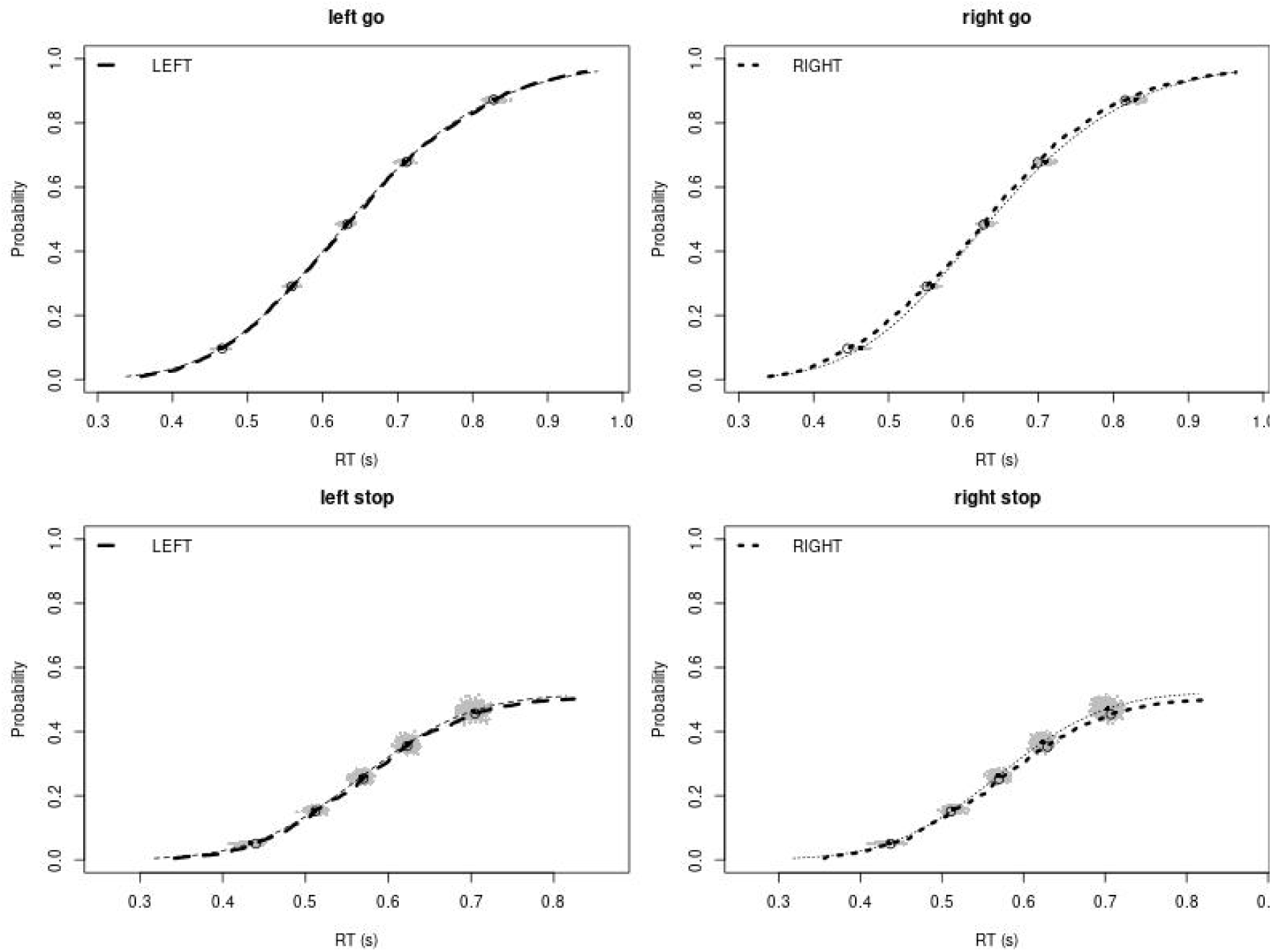
Observed and predicted cumulative distribution function (CDF) of Go-RTs (upper panels) and signal-respond RTs (lower panels), separately for left and right stimuli, for the non-rIFC lesion group after removing stop-signal trials at SSDs of 0, 50, 100, 150, and 250 ms. The observed and predicted CDFs were averaged across participants. Signal-respond RTs were collapsed across SSD. Thick dashed and dotted lines show the CDF of the observed “LEFT” and “RIGHT” responses, respectively. Circles show the 10^th^, 30^th^, 50^th^, 70^th^, and 90^th^ percentile of the distributions. Thin dashed and dotted lines show the CDF of the predicted “LEFT” and “RIGHT” responses, respectively, averaged across the 500 predictions. For each percentile, the gray clouds show the 500 predicted percentiles.

**Figure S10.**
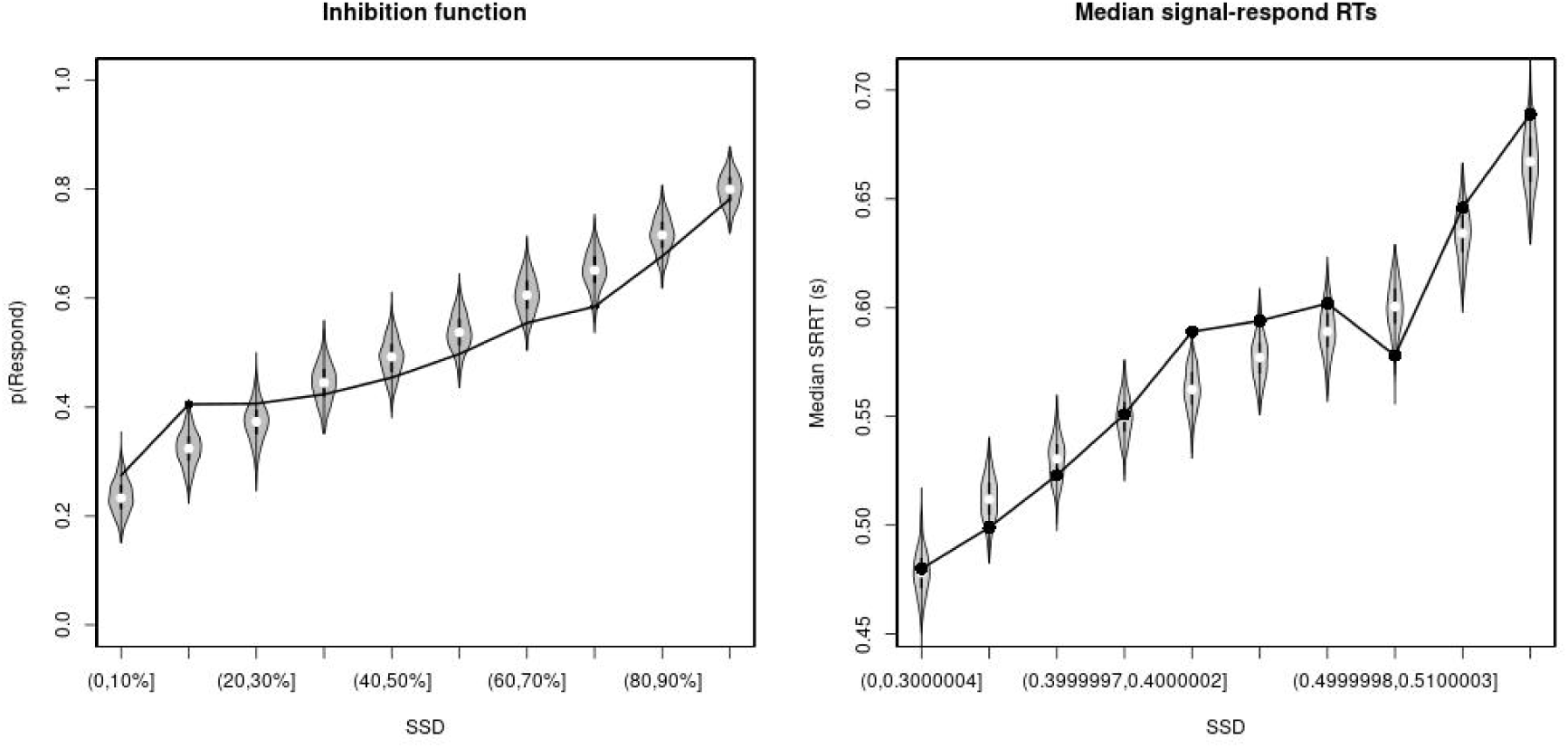
Observed and predicted inhibition function (left panel) and median signal-respond RT as a function of stop-signal delay (SSD; right panel) for the non-rIFC lesion patients after removing stop-signal trials at SSDs of 0, 50, 100, 150, and 250 ms. In the left panel, black bullets show the observed average response rate on stop-signal trials (P(Respond)) for each SSD category, where the SSD categories were defined in terms of the percentiles of the distribution of SSDs for each participant and then averaged across participants. In the right panel, black bullets show the observed average median signal-respond RT (SRRT) for each SSD category, where SSD categories were defined by pooling SSDs over participants before calculating the percentiles. The gray violin plots show the distribution of the 500 average response rates and SRRTs predicted by the model, with the white circles representing the median of the predictions.

**Figure S11.**
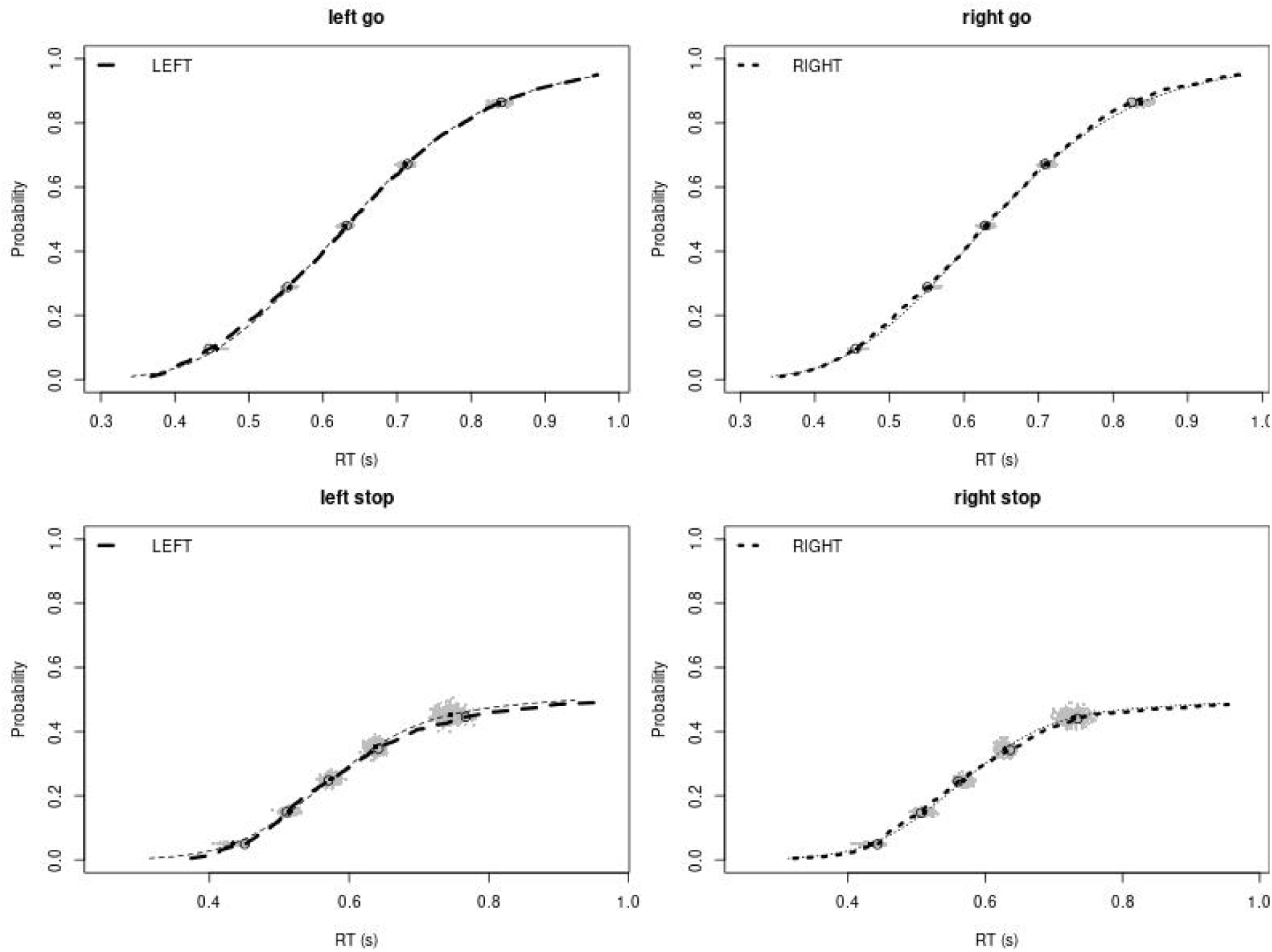
Observed and predicted cumulative distribution function (CDF) of Go-RTs (upper panels) and signal-respond RTs (lower panels), separately for left and right stimuli, for the matched comparison group for the non-rIFC lesion patients after removing stop-signal trials at SSDs of 0, 50, 100, 150, and 250 ms. The observed and predicted CDFs were averaged across participants. Signal-respond RTs were collapsed across SSD. Thick dashed and dotted lines show the CDF of the observed “LEFT” and “RIGHT” responses, respectively. Circles show the 10^th^, 30^th^, 50^th^, 70^th^, and 90^th^ percentile of the distributions. Thin dashed and dotted lines show the CDF of the predicted “LEFT” and “RIGHT” responses, respectively, averaged across the 500 predictions. For each percentile, the gray clouds show the 500 predicted percentiles.

**Figure S12.**
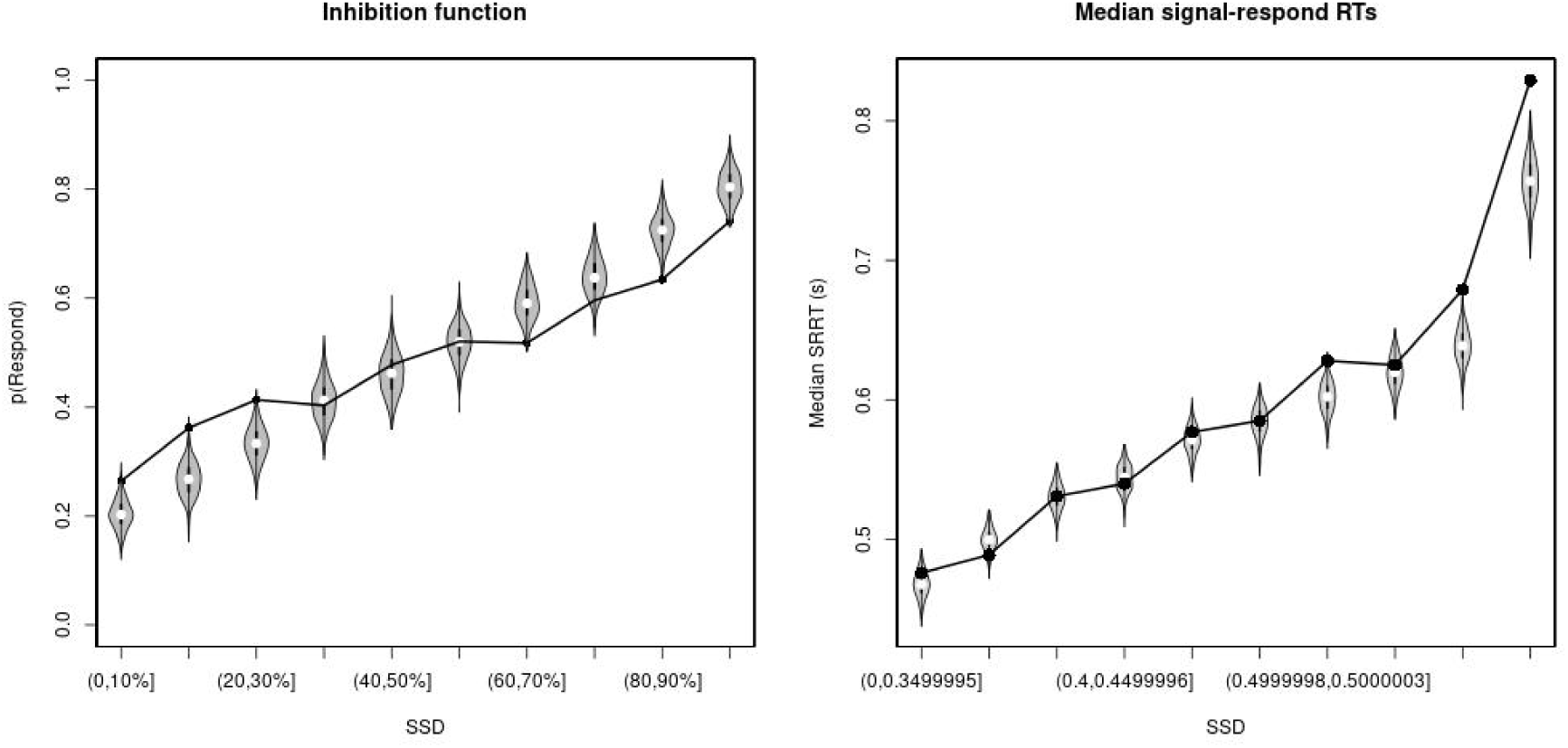
Observed and predicted inhibition function (left panel) and median signal-respond RT as a function of stop-signal delay (SSD; right panel) for the matched comparison group for the non-rIFC lesion patients after removing stop-signal trials at SSDs of 0, 50, 100, 150, and 250 ms. In the left panel, black bullets show the observed average response rate on stop-signal trials (P(Respond)) for each SSD category, where the SSD categories were defined in terms of the percentiles of the distribution of SSDs for each participant and then averaged across participants. In the right panel, black bullets show the observed average median signal-respond RT (SRRT) for each SSD category, where SSD categories were defined by pooling SSDs over participants before calculating the percentiles. The gray violin plots show the distribution of the 500 average response rates and SRRTs predicted by the model, with the white circles representing the median of the predictions.

